# Harnessing hydrodynamics for high-yield production of extracellular vesicles from stem cells spheroids with specific cargo profiling

**DOI:** 10.1101/2025.01.02.631101

**Authors:** Solène Lenoir, Elliot Thouvenot, Giacomo Gropplero, Léonie Dec, Damarys Loew, Clotilde Théry, Jose E Perez, Claire Wilhelm

## Abstract

This study presents a novel method and device for the hydrodynamic production of extracellular vesicles (EVs) derived from biomimetic multicellular 3D spheroids, enabling high-throughput particle release that is 10 to 20 times higher than in non-stimulated conditions. The device facilitates the formation of spheroids from human mesenchymal stem cells (hMSCs), offering an all-in-one approach for both spheroid generation and EV release. Production times are reduced to just a few hours, with yield further increased by alternating periods of high hydrodynamic flow and spheroid recovery in a sequential production approach. Using this system, we explored the impact of hydrodynamic and starvation conditions on the protein cargo of EVs, identifying distinct protein markers through proteomics. Specifically, hydrodynamic stimulation enriched EVs in plasma membrane-derived and mitochondrial proteins, revealing divergent biogenesis pathways. Importantly, the produced EVs exhibited therapeutic properties, with demonstrated effects in wound healing, angiogenesis, and anti-inflammatory responses, some showing enhanced efficacy under hydrodynamic stimulation.

## INTRODUCTION

Extracellular vesicles (EVs)^1^, particularly those derived from mesenchymal stem cells (MSCs)^2^, have emerged as potential therapeutic agents in regenerative medicine^3^. These membrane-bound vectors, ranging from 50 to 300 nm, were initially identified for their role in intercellular communication but are now recognized for carrying the therapeutic properties of their parent cells. Besides, the therapeutic effects of stem cells themselves are increasingly attributed to paracrine mechanisms, primarily mediated by EVs^4^. This transfer of therapeutic functions from MSCs to EVs provides a cell-free alternative to traditional cell therapy, minimizing the risks associated with living cell treatments, such as unwanted replication and differentiation^5^. Another notable advantage of EVs is their immune-privileged status, which facilitates allogeneic use as an off-the-shelf therapeutic option. Additionally, the use of EVs is further enhanced by their improved storage capabilities and shelf-life compared to whole cells^6^.

MSC-derived EVs are now recognized for their ability to restore tissue function and homeostasis by modulating cellular processes, as demonstrated in the treatment of injuries to the heart, kidney, liver, brain, or skin^7–14^. They are involved in every stage of the regenerative process^15–17^, facilitating cell migration and angiogenesis, influencing proliferation, managing cell senescence and differentiation, or reducing heart ischemia-reperfusion injury^18–21^. Their potential in cancer therapy is also compelling, as they have demonstrated the ability to inhibit proliferation and resistance to apoptosis^22^.

Given these significant therapeutic perspectives on EVs and their potential for widespread application, one might expect them to already be in clinical use as a replacement for cell therapies. However, the transition of EV-based therapies from research to clinical practice still faces several challenges, in engineering^23–25^, production^26, 27^, isolation^28–32^, storage^33^, standardization and commercialisation^34^.

Concerning production, the primary challenge is to increase the yield of EV production. Various approaches are employed for this purpose, utilizing different modes of stimulation. A widely adopted approach for enhancing EV production involves culturing cells under stress conditions, such as hypoxia, nutrient deprivation or temperature, which have been shown to boost EV output^35–42^. Biochemical techniques that incorporate specific growth factors, cytokines, or small molecules have also been effective in promoting the release of EVs^43^. Genetic modification of donor cells to upregulate the expression of EV-related proteins is another relevant strategy. For instance, overexpression of the tetraspanin CD9 results in a significant increase of EV production^44^. For these approaches to provide sufficient EVs for clinical use, large-scale cell culture platforms, such as hyperflasks or 3D cultures on microcarriers, are necessary to increase the number of cells producing EVs. While these methods perform well^45^, they remain both material and time consuming. Recently, the use of physical stimuli, such as shear stress or mechanical agitation, has been introduced to effectively trigger the secretion of EVs from donor cells^46–49^. This approach is founded on the shear stress present in our body, such as in blood vessels^50–52^ or during cardiac pumping^53^. Initial attempts to trigger EV release through shear stress involved using microfluidic devices with micro-channels smaller than cells^52, 54, 55^. While these methods enabled rapid EV release, they still faced challenges related to yield and scalability. To increase yield, culturing cells in hollow-fiber bioreactors is an excellent option^56, 57^, although it requires prolonged production times, resulting in significant delays in generating sufficient EV quantities for clinical use.

The goal of this study is to address these limitations by proposing an alternative hydrodynamic-based strategy to enhance EV yield and scalability. The rationale is to leverage dynamic hydrodynamic cell cultures operating in a high shear turbulent regime to stimulate EV release. It demonstrates a significant enhancement of EV release using a specially designed and manufactured device that incorporates an alternating rotation tube with internal obstacles. Beyond the innovative and efficient bioproduction technology, the uniqueness of this strategy lies in the release of extracellular vesicles from human stem cells (hMSCs) organized as spheroids that mimic their native 3D tissue-like environment, that can modulate EVs properties ^58–60^. Additionally, the device was enhanced to streamline both the formation of spheroids and the massive release of EVs from the resulting spheroids. To facilitate the adoption of these hydro-spheroid-produced EVs and fully realize their potential as a cost-effective, cell-free solution, it was essential to assess their cargo and potency. This evaluation included whole proteome analysis, along with functional tests for wound healing, angiogenesis and anti-inflammatory properties. Hydro-spheroid-produced EVs exhibited the typical markers associated with EVs, but they also revealed a distinct hydrodynamic and spheroid-derived cargo signature when compared to those produced through starvation, and notably demonstrated enhanced functional properties.

## RESULTS

### A rapid and efficient hydrodynamic method enhances EV bioproduction in hMSC spheroids

The objective was to design a device to explore ways of controlling hydrodynamic shear flow-mediated bioproduction of EVs from human stem cells organized in a spheroid 3D biomimetic environment. Such process involves creating a turbulent flow with vortices that trap spheroids, subjecting them to high hydrodynamic stresses, that stimulate the production of spheroids-derived EVs. The method, patented in 2023, employs rotating tubes featuring turbulence-enhancing inner obstacles and operating in a reversing mode (Fig. 1a), to introduce an additional layer of shear flow stresses. The design of the tube is fully modular, including the internal obstacles. Two configurations were tested and 3D printed, including inner walls and grids structures (Supplementary fig. 1). Both performed efficiently (Supplementary fig. 2), so the baffled design (with inner walls) was selected as one that could be manufactured from plastic by injection molding, offering the advantages of high-throughput production, easy sterilization, and optimal biocompatibility. It features three pairs of inner walls oriented at 120° to each other, with heights increasing from 1 cm to 2 cm to 4 cm, in a tube with a diameter of 3.7 cm and a total height of 8.4 cm. The tube’s geometry enables real-time tracking of flow dynamics, resolving of the turbulent regime by particle image velocimetry (Fig. 1b). Velocities as high as 10 cm/s were reached, corresponding to Reynolds number in the 2000-20000 range.

With this specific design, the tube holds between 20 and 30 mL, typically 25 mL, which corresponds to a two-thirds fill level. It is filled with medium containing the cellular material, prepared strictly without serum for the production stages. Fig. 1c illustrates the interior of the tube when it contains spheroids that were previously formed through micropatterning (Fig. 1d). The spheroids were generated in 200 µm agarose-made microwells within large wells of a 6-well plate, featuring 2,000 microwells per well. A total of 500,000 hMSCs were used per well, and the cells matured over 2 days to form dense spheroids (Supplementary fig. 3). The spheroids were then resuspended in serum-free medium at a concentration of 2,000 spheroids per mL (comprising all spheroids from a single well in one mL), enabling one tube to hold the equivalent of the content from four entire 6-well plates. As proof of concept, Fig. 1e shows the number of particles released per cell (accounting for all cells within the spheroids) during stimulation for either 1 hour or 3 hours at 800 rotations per minute (rpm), with a direction reversal period of 5 seconds. This production is compared with the measurements obtained after 3 hours under static conditions (the same spheroids in a non-adherent flask at the same density), resulting in an almost 20-fold increase due to hydrodynamic stimulation. This measurement of the number of particles produced was conducted using nanoparticle tracking analysis (NTA) techniques, utilizing either the Videodrop (Myriade) instrument or the NanoSight (Malvern) instrument. The NanoSight has a lower detection limit of 50 nm, compared to the Videodrop, which has a limit of 80 nm. This accounts for the greater number of particles measured by the NanoSight. Furthermore, Supplementary fig. 4 demonstrates linear correlation across all measurements conducted in this study on the same samples using both instruments, with an average of 4.2 times more particles measured by the NanoSight. To compare with other bioproduction techniques, the values provided by the NanoSight, which is the instrument mostly used, must be considered. For rotation condition (ROT), more than 30,000 particles are produced per cell, placing it at the high end of bioproduction methods^61^. For example, in hyperflasks, bone marrow BM-MSCs produce 6,000 EVs per cell over 48 hours^62^, as measured by NanoSight. The intermittent rotation baffled tube technology thus produces 5 times more EVs in one-tenth the time compared to hyperflasks. When compared to flow-based production, the state-of-the-art in bioproduction is certainly the hollow fiber system. Using UC-MSCs, hollow fiber systems produce, assuming correct calculations, 50 EVs per cell (NanoSight measurement) per 3-day harvest^63^, with cultures maintained for 55 days, resulting in 18 harvests, and ultimately 1,000 EVs per cell in 1320 hours. This indicates a production rate 30 times higher in 400 times less time. However, the scale-up of hollow fibers has been achieved with Quantum bioreactors, which, with BM-MSCs, produce 15,000 EVs per cell (NanoSight measurement)^64, 65^, closer to the order of magnitude of the rotating tubes, but over 4 to 21 days, versus 3 hours.

Figures 1F to 1H focus on the direct impact of hydrodynamic stimulation on the viability of spheroids, the initial mandatory step in evaluating an EV bioproduction technology. Metabolic activity was measured using resazurin metabolic conversion to resorufin (Alamar Blue) after 1 hour or 3 hours of production under static control conditions and under rotation condition (800 rpm), expressed as a percentage of the initial activity. The results showed a slight decrease in metabolic activity after 3 hours, approximately 20%. Concurrently, cell death was assessed by measuring the release of the enzyme adenylate kinase from damaged cells (ToxiLight). There was a good correlation with metabolic activity, showing around 20% cell mortality. These findings were corroborated by a live/dead assay at the spheroid level (Figures 1G-1H), which indicated a slight decrease in green calcein signal (intracellular esterase activity of live cells) and a corresponding increase in red ethidium signal (loss of plasma membrane integrity in dead cells) only after 3 hours of production.

All subsequent measurements will be reported using Videodrop to assess yield and ToxiLight to assess cell death. As just demonstrated, both methods were validated relative to other techniques. They provide the only viable procedure for evaluating numerous stimulation parameters of the bioproduction process. Indeed, Videodrop allows for much quicker measurements compared to NanoSight (30 seconds versus 5 minutes), requiring only 8 µL of the conditioned medium (CM) per measure, while ToxiLight can also be performed on the CMs, using only 5 µL per measure. Thus, Fig. 1i illustrates the particle production per cell measured by Videodrop as a function of production time and intermittent rotation speed, while Fig.1j shows the same conditions but in terms of mortality assessed by ToxiLight. There is a kinetic effect, with production increasing linearly during the first hour before gradually saturating, whereas mortality increases linearly with production time, becoming significant at 1600 rpm.

The final step in the proof of concept for the production method is to initiate production directly with suspended individual cells, bypassing the spheroid formation stage in the microwells. This corresponds to Fig. 1k (particles per cell) and Fig. 1l (mortality), which demonstrate similar viability—albeit slightly lower—and, most importantly, comparable production rates, while in this condition with individual cells, all cells are accessible to the flow. Spheroids thus produced (per cell) as much as single cells, where one might have thought that the core was non-productive.

Finally, the particles produced were observed by cryo-electron microscopy to provide detailed visualization of vesicular morphology. The conditioned media were first concentrated and purified using size-exclusion chromatography (SEC), which effectively isolated EVs from soluble factors before electron microscopy observations. Control EVs were produced and observed from starvation stimulation conditions (0 rpm, 72 hours) with individual cells in a monolayer (2D, Fig. 1m) or as spheroids (3D, Fig. 1n). These were compared with EVs from hydrodynamic stimulations at 600 rpm with individual cells (Fig. 1o) and spheroids (Fig. 1p). Supplementary images for these conditions are provided in Supplementary figs. 5, 6, 7 and 8, with more than 40 individual EVs observed per condition. All vesicles observed were intact, and their diameters were measured (Fig. 1q), revealing comparable size distributions with an average diameter of 85 nm.

**Fig. 1:**
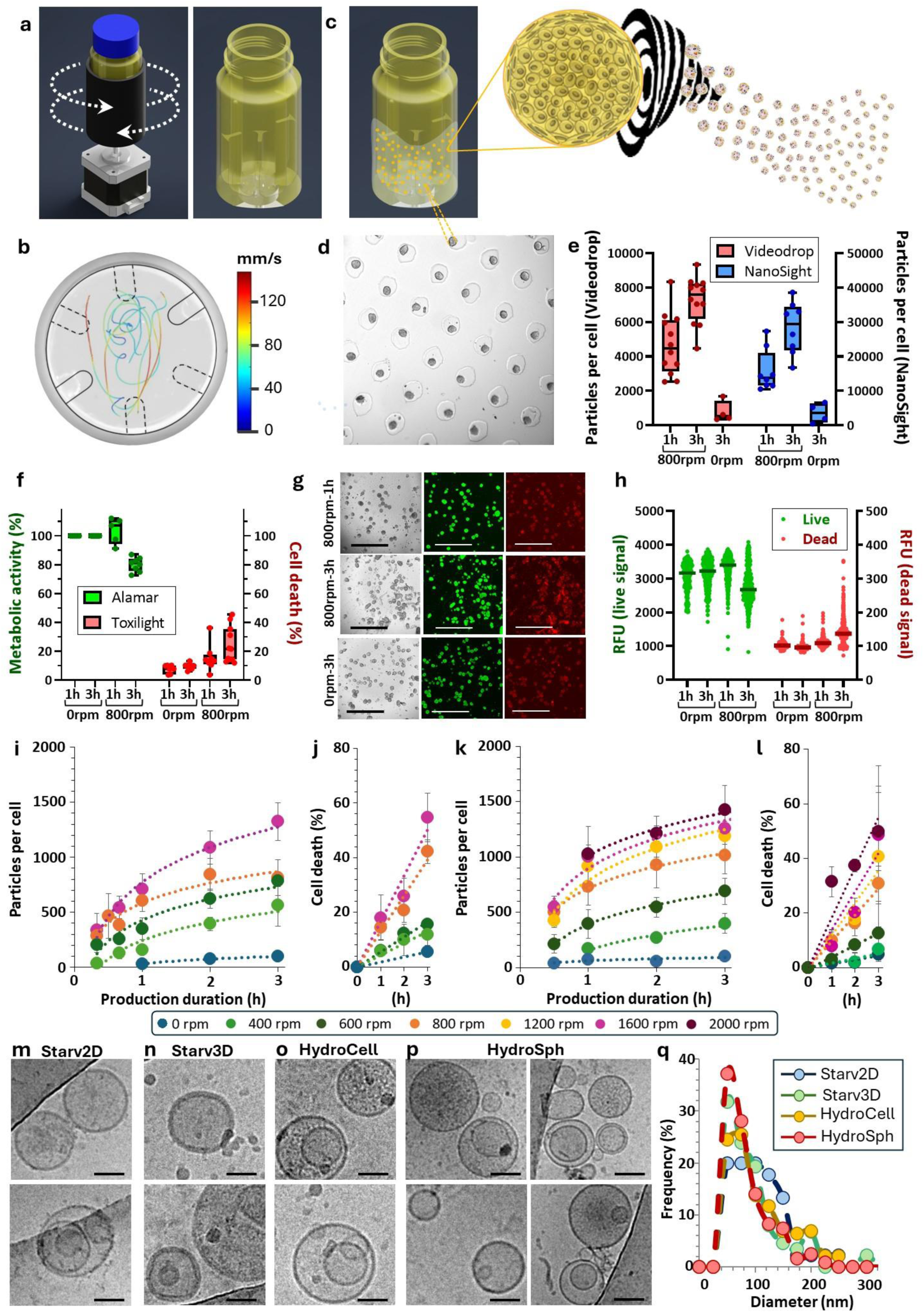
Bioproduction of EVs from hMSCs in rotating tubes. **a**: Our new technology: back-and-forth rotating tubes with interior broken walls that enhance turbulence within. The tube rotates, stops, and then reverses direction. **b**: Tracking the movement of phosphorescent beads in a plane of the rotating tube (800 rpm) allows for direct visualization of the turbulent flow. **c**: Schematic of spheroids trapped in the flow, with vortex formation generating intense flow stresses on each spheroid, leading to the production of many EVs. **d:** Bright field imaging of the agarose microwell array that allows for spheroid formation before injection into the tubes. Each spheroid forms in an individual microwell, starting from 250 individual cells, resulting in spheroids with an average diameter of 125 ± 20 µm. **e:** Measurements of the number of particles produced per cell for 1 hour or 3 hours of hydrodynamic stimulation (800 rpm) or 3 hours under static conditions (0 rpm). Each point represents an independent biological replicate, and each condition was measured by NTA using both the Videodrop and NanoSight instruments. **f:** After 1 hour or 3 hours of production under hydrodynamic (800 rpm) or static (0 rpm) conditions, each condition was measured using ToxiLight to assess cell mortality and AlamarBlue to evaluate metabolic activity. Each point represents an independent biological replicate. **g:** Live (green) and dead (red) imaging of spheroids after 1 hour or 3 hours at 800 rpm or 0 rpm. **h:** Quantification of the live and dead signal for the different conditions, with each point corresponding to measurements on an individual spheroid. **i, j:** Kinetic curves of particle production per cell (organized in spheroids) measured by Videodrop (I) and spheroid mortality measured by ToxiLight (J) for six conditions: 0, 200, 400, 800, 1200, 1600, and 2000 rpm. Each point represents the mean, and the error bars indicate standard deviation, with n > 3 for all conditions. **k, l:** Similar curves of particle production per cell (K) and mortality (L) as a function of production time, but for production from individual cells in suspension, not yet organized into spheroids, with n > 3 for all conditions. **m-p**: Cryogenic transmission electron microscopy imaging of extracellular vesicles produced from spheroids (600 rpm Sph, P) or individual cells (600 rpm Cell, O) in hydrodynamic conditions, and for EVs produced upon serum starvation from spheroids (0 rpm 3D, N) or adherent cells (0 rpm 2D, M). **q**: Diameter distributions extracted from cryoTEM imaging (n=45 for Starv2D, n=87 for Starv3D, n=120 for HydroSph, n=190 for HydroCell).

### Proteomic analysis reveals that EVs generated under hydrodynamic flow exhibit traditional EV markers but that their protein content only partially overlaps with starvation-derived EVs

We investigated the protein content of EVs from four different conditions—Starv2D, Starv3D, HydroCell, and HydroSph—each produced as five biologically independent replicates, using label-free mass spectrometry-based proteomics. The Starv2D condition refers to the classical control of 72 hours of starvation in adherent 2D cultures (0 rpm – 2D). The Starv3D condition involves spheroids placed on non-adherent substrates, identical to those used in rotation stimulation, also subjected to 72 hours of starvation (0 rpm – 3D). The HydroCell and HydroSph conditions involve 3 hours of hydrodynamic stimulation at 600 rpm, with HydroCell starting from individual cells in suspension and HydroSph from spheroids. Additionally, we included as a positive control for hydrodynamic conditions a millifluidic hydrodynamic stimulation of spheroids in a cross-slot chip^66^ that creates a single vortex with an imposed Reynolds number of 400 (referred to as HydroChip). The protein content was measured for the five independent replicates, with values expressed as the number of particles per µg of protein. All values were found to be in the range of 10^8^, specifically (0.9±0.2)x10^8^ for Starv2D, (1.1±0.6) x10^8^ for Starv3D, (1±0.3) x10^8^ for HydroCell, (1.4±0.6) x10^8^ for HydroChip and (1.1±0.3) x10^8^ for HydroSph.

A total of 6688 proteins with at least 2 distinct peptides were identified, with 6125 detected in Starv2D, 6042 in Starv3D, 6595 in HydroCell, 6589 in HydroSph, and 6532 for HydroChip. Among these, 4638 proteins were common across all samples. Principal component analysis (PCA) was used to examine the differences in protein composition among the groups (Fig.2a). This unbiased method clearly distinguished different EV populations, separating the hydrodynamic conditions (with the HydroChip condition clustering with HydroCell and HydroSph) from the starvation conditions and further distinguishing between 2D and 3D environments. Hierarchical clustering analysis based on normalized protein intensities is represented for all proteins and samples in Supplementary fig. 10). Gene ontology enrichment analysis was performed for the protein lists of each cluster compared to the entire human proteome. The four Gene Ontology terms identified in this study were extracellular vesicles (1133 proteins), wound healing (156 proteins), angiogenesis (124 proteins), and mitochondria (730 proteins). These proteins are presented in the volcano plots of Fig. 2, alongside all proteins across all replicates. For these volcano plots, proteins were selected based on the presence of at least two distinct peptides in at least one replicate across all conditions, and an adjusted p-value of less than 0.05. The conditions were compared pairwise: the Starv2D condition, with individual adherent monolayer cells under starvation, was compared with the HydroCell condition, with individual cells under hydrodynamic stimulation (Fig. 2b, 2d, 2fF); and the Starv3D condition, with spheroids under starvation, was compared with the HydroSph condition, with spheroids under hydrodynamic stimulation (Fig. 2c, 2e, 2g). Proteins with a log2(fold change) between -2 and 2 were considered common to the two conditions. For the HydroCell and HydroSph conditions, enrichment was considered significant for a log2(fold change) > 2, while for Starv2D and Starv3D conditions, it was considered significant for a log2(fold change) < -2. For the term extracellular vesicles, there was greater enrichment in the hydrodynamic conditions, with 13% (149 proteins) enriched in HydroCell versus 10% (117 proteins) enriched in Starv2D; and 19% (215 proteins) enriched in HydroSph versus 6% (68 proteins) in Starv3D. Wound healing and angiogenesis showed similar enrichment between hydrodynamic and starvation conditions. For wound healing, 6% (10 proteins) versus 15% (24 proteins) were enriched in HydroCell versus Starv2D; and 12% (18 proteins) versus 9% (13 proteins) were enriched in HydroSph versus Starv3D. For angiogenesis, 10% (12 proteins) versus 18% (23 proteins) were enriched in HydroCell versus Starv2D; and 12% (15 proteins) versus 9% (11 proteins) were enriched in HydroSph versus Starv3D. In contrast, a significant mitochondrial signature was identified in hydrodynamic conditions, for both 2D and 3D comparisons: 32% (234 proteins) versus 0.6% (4 proteins) were enriched in HydroCell versus Starv2D; and 31% (221 proteins) versus 0.4% (3 proteins) were enriched in HydroSph versus Starv3D.

For the Z-score heatmaps shown in Figures 2h to 2l, all four conditions were included, and proteins were selected with at least 2 distinct peptides across all 5 biological replicates of an experimental condition. The proteins are listed in the heatmaps. For the terms wound healing (Fig. 2j), angiogenesis (Fig. 2k), and mitochondrion (Fig. 2l), this narrowed the number of proteins to 36, 23, and 101, respectively. For wound healing and angiogenesis, as predicted by the volcano plots, the distribution of proteins was relatively uniform across all conditions. In contrast, for mitochondria-associated proteins, the Z-score was significantly higher for most proteins in the hydrodynamic stimulation conditions, again consistent with the overexpression observed in the volcano plots. For the term extracellular vesicles, the heatmaps were restricted to EV markers as outlined in the Minimal Information for Studies of Extracellular Vesicles (MISEV2023) guidelines^67^. Of the 60 proposed transmembrane markers (MISEV category 1, Fig. 2h) and 47 cytosolic markers (MISEV category 2, Fig. 2i), 42 and 28, respectively, were detected across the different conditions of the proteome analysis. The co-presence of both transmembrane and cytosolic markers indicates the presence of intact vesicles, as recommended by the MISEV guidelines. Western blot analysis (Supplementary fig. 11) confirmed the presence of transmembrane domain proteins CD63 and CD81, as well as the cytosolic protein Syntenin-1, in all conditions, while common contaminants such as 14-3-3 proteins were absent. Similarly, MACSplex analysis^68^ (Supplementary fig. 12) verified the presence of CD63 and CD81, along with mesenchymal-associated markers, CD29, CD44, CD49e, CD105, and CD146, across all conditions.

The transmembrane markers (Fig. 2h) are distributed without a distinct pattern among the different conditions, whereas the Z-scores of detected cytosolic markers (Fig. 2i) indicate distinct biogenesis pathways for the two types of stimulation. EVs generated under starvation conditions are markedly enriched in exosome-associated proteins, while EVs produced under hydrodynamic conditions exhibit significant enrichment in microvesicle-related proteins. To further investigate, we identified the contribution of the ESCRT machinery (Fig. 2m). The four ESCRT complexes and the majority of proteins constituting the ESCRT system (except EPS15, an accessory protein of ESCRT-0, and CHMP6 (VPS20) in ESCRT-III) are overexpressed in the static starvation condition compared to the hydrodynamic condition. Most proteins involved in ESCRT-independent exosome mechanisms (including CD63) are underexpressed in the hydrodynamic condition. Additionally, ARF6, involved in ESCRT-independent microvesicle formation, and all the flippases are overexpressed under hydrodynamic conditions. This suggests that EVs formed under hydrodynamic conditions are produced through an ESCRT-independent mechanism, likely via plasma membrane budding.

Overall, these proteomics proteomics data reveal that hydrodynamic stimulation induces a distinct proteome signature in the EVs produced, characterized by the enrichment of microvesicle-related proteins, and primarily formed through an ESCRT-independent pathway. Therefore, the hydrodynamic stimulation not only enhance EV production but also induces a distinctive proteomic profile, setting them apart from those produced under static conditions. The mitochondrial signature, highlighted by the enrichment of mitochondrial-associated proteins, suggests a role in the energy metabolism and biogenesis processes of the EVs, also confirmed by the ESCRT-independent mechanism. Such a tailored proteomic signature could be advantageous for specific therapeutic applications where enhanced metabolic activity or mitochondrial function is desired. Furthermore, this approach underscores the versatility and precision of hydrodynamic stimulation in modulating the content of EVs, for the development of specialized EV-based therapies.

**Fig. 2:**
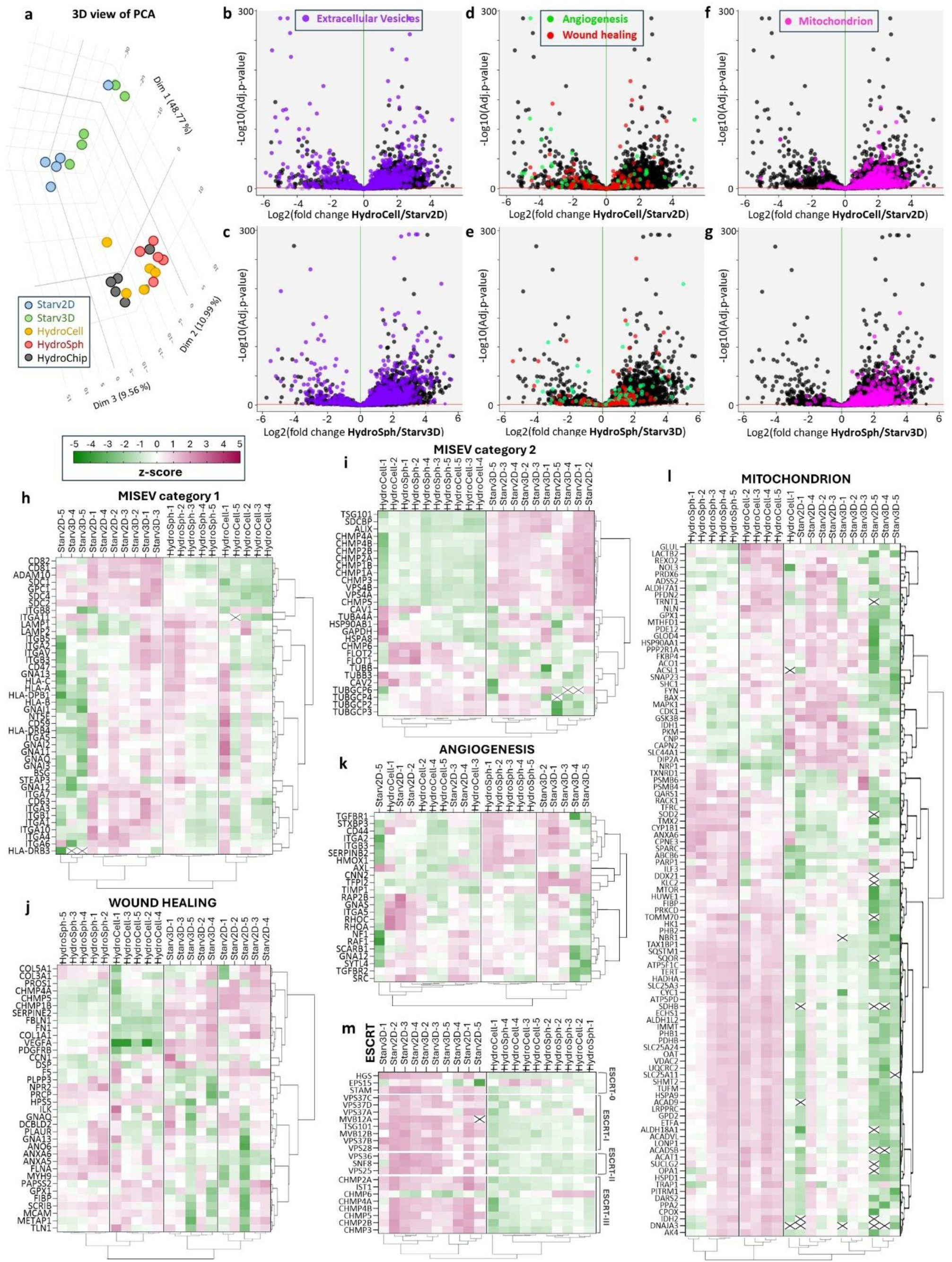
Proteome characterization by mass spectrometry-based proteomics. **a**: 3D view of Principal Component Analysis (PCA) of the proteome for conditions Starv2D (blue), Starv3D (green), HydroCell (yellow), HydroSph (red) and HydroChip (black). **b-g**: Proteome comparison between conditions depicted in volcano plots. Proteins associated with the “extracellular vesicle” GO term (GO:1903561) are shown in purple, on top of the total of proteins quantified in both conditions (black). Comparisons are made between Starv2D and HydroCell (**b**), and Starv3D and HydroSph (**c**). Proteins associated with the “wound healing” GO term (GO:0042060) and “angiogenesis” GO term (GO:0001525) are represented in red and green, respectively, and are shown for comparisons between Starv2D and HydroCell (**d**), or Starv3D and HydroSph (**e**). Proteins associated with the “mitochondrion” GO term (GO:0005739) are shown in pink for comparisons between Starv2D and HydroCells (**f**), and Starv3D and HydroSph (**g**). **h-l**: Heatmaps with hierarchical clustering of EVs produced through serum starvation from hMSC in 2D configuration (Starv2D) and 3D configuration (Starv3D) and EVs produced from individual hMSCs (HydroCell) or hMSC-derived spheroids (HydroSph) stimulated hydrodynamically in the reverse rotating tubes. The heatmaps display Z-scores for all proteins found, including transmembrane EV markers as described in the MISEV category 1 (**h**), cytosolic EV markers as described in the MISEV category 2 (**I**), for proteins related to wound healing (**j**), angiogenesis (**k**), mitochondria (**l**), and ESCRT-related proteins (**m**).

### Hydrodynamic stimulation increases EVs potency

The next step was to test the EVs’ functions in wound healing69, angiogenesis20, and anti-inflammatory activity.

We used a scratch test to measure the EVs’ wound healing ability to induce fibroblast migration by testing EVs on human dermal fibroblasts (HDFs) seeded in 96-well plates, with an artificial wound created by scratching a 1 mm gap at the center of each well. After the scratch, EVs and serum were administered, and wound closure was monitored over time by live imaging at 24, 48, and 72 hours (Fig. 3a, other images in Supplementary fig. 13A-H). For the EVs (produced under the conditions Starv2D, Starv3D, HydroCell, and HydroSph), the dose used was 10^9^ EVs per mL, equivalent to 10^8^ EVs per wound. These conditions were compared to medium alone and medium supplemented with FBS at concentrations of 2%, 4%, and 10%. FBS addition gradually increased wound closure capacity. At all time points, the four EV conditions exhibited significantly higher reparative potential compared to FBS0% and FBS2%. The Starv2D condition yielded results similar to FBS4% but lower than FBS10% at 24 and 72 hours. The other three EV conditions produced outcomes comparable to FBS10% at all time points, with a non-significant trend towards improved wound healing, except for the HydroSph condition at 72 hours, which showed a significant increase. Overall, EVs produced from 3D spheroids, either by starvation or by hydrodynamic stimulation, performed better in improving wound colonization compared to FBS at 4%, were similar to FBS at 10%, and were even more effective at 72 hours for the hydrodynamic stimulation. Hydrodynamically stimulated EVs thus significantly enhanced the wound healing process, surpassing the effect of serum proteins at 1-4%, exhibiting a comparable effect to serum proteins at 10%, and demonstrating slightly greater efficacy than EVs produced using the classical method of starvation.

Next, the neo-angiogenic potential of EVs was evaluated using human umbilical vein endothelial cells (HUVECs) cultured as 3D spheroids in agarose microwells and incorporated into a collagen matrix (Fig. 3a). Similar conditions to those used for the wound healing assay were applied, testing only FBS0% (negative control) and FBS10% (positive control), with an additional positive control of VEGF administration at different doses (12.5, 25, and 50 ng/mL). Treatment of these spheroids with VEGF or serum proteins (FBS) induced sprout formation (Fig. 3c, additional images in Supplementary fig. 14). Quantification of the number of sprouts is shown in Fig. 3d, and Supplementary fig. 15 presents the sprout lengths. The administration of 5 x 10^7^ (10^9^ per mL) and 10^8^ EVs (2 x 10^9^ per mL) per endothelial spheroid gradually increased the number of sprouts per spheroid across all EV conditions, indicating their significant role in promoting angiogenesis.

Finally, the anti-inflammatory potential of EVs was evaluated using RAW264.7 macrophages treated with lipopolysaccharide (LPS) to induce nitric oxide (NO) production, an indicator of inflammation. EVs generated under hydrodynamic conditions, namely HydroCell and HydroSph, demonstrated a significant, dose-dependent reduction in NO formation. At the highest concentration tested (4x = 3 x 10^8^), these hydrodynamically produced EVs nearly completely suppressed NO production, achieving an effect comparable to that of dexamethasone 8 µg/mL (8x), thereby showcasing their potent anti-inflammatory properties. We hypothesize that this notable anti-inflammatory effect is associated with the overexpression of mitochondria-related proteins in the hydrodynamically produced EVs. LPS-induced inflammation in macrophages is known to result in mitochondrial damage and compromised mitochondrial activity and integrity^70^. Restoring mitochondrial integrity and promoting biogenesis are crucial for mitigating and reversing inflammation, as well as preventing excessive responses to LPS stimulation^71^. Emerging research has indicated that EVs can transport and deliver mitochondrial contents, including mitochondrial DNA (mtDNA) and proteins to recipient cells^72^. Notably, recent studies have shown that EV-mediated transfer of mitochondrial components can enhance mitochondrial integrity and function in recipient cells, thereby reducing inflammation^73^ and promoting an anti-inflammatory phenotype in LPS-stimulated macrophages^71^. Thus, we propose that the observed anti-inflammatory effect of hydrodynamically produced EVs is due to their ability to restore mitochondrial integrity by delivering mitochondria-related proteins to LPS-stimulated macrophages. This mechanism likely underpins the significant reduction in NO production and the overall anti-inflammatory activity observed in our study.

**Fig. 3:**
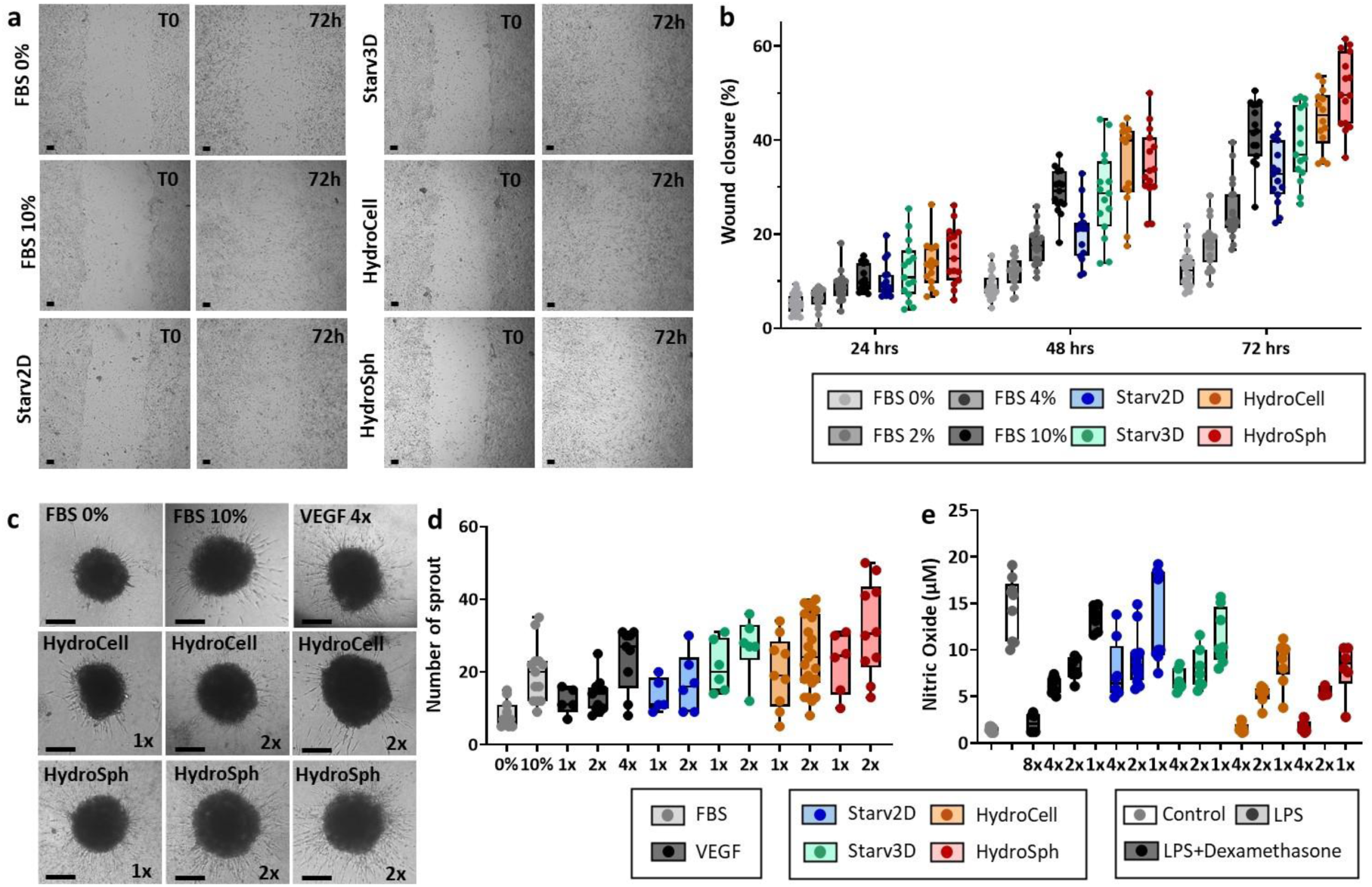
Functionality tests. EVs produced by static starvation in monolayers (Starv2D) or spheroids (Starv3D), as well as EVs produced by hydrodynamic stimulation in reverse rotating baffled tubes from either individual cells (HydroCell) or spheroids (HydroSph), were tested for wound healing (**a-b**), neo-angiogenesis (**c-d**), and anti-inflammatory activity (**e**). **a**: For wound healing, representative images depicting the initial wound gap (left) and wound healing at the 72-hour time point (right) are provided for all conditions. Scale bar = 100 µm. Positive controls for this test are serum conditions with different volume fractions (1, 2, 4, and 10%), and the negative control corresponds to medium alone (no EVs, and 0% FBS). The administered dose (concentration measured with Videodrop) of EVs is 1x10^9^ EVs per mL (i.e. 10^8^ EV per wound). **b**: Time course of the effect of serum proteins or different EVs on wound healing over 4 days. **c**: Typical images of sprout formation from endothelial spheroids, illustrating the neo-angiogenesis capacity. Scale bar = 100 µm. Various conditions were tested, including serum proteins (FBS) diluted to 10% (FBS 10%), no addition (FBS 0%), VEGF at three doses 12.5 (1x), 25 (2x), 50 (4x) ng/mL, and EVs from the conditions Starv2D, Starv3D, HydroCell, and HydroSph (1x = 5x10^7^ EVs, equivalent to a concentration of 10^9^ EVs per mL). **d**: Number of sprouts measured for all conditions. **e**: Testing anti-inflammatory activity of EVs produced from all conditions on human macrophages (RAW264.7) treated with lipopolysaccharide (LPS). LPS induced the formation of nitric oxide (NO), indicating an inflammatory reaction. Dexamethasone at different doses (1x = 1µg/mL) and EVs from all conditions (1x = 7.5x10^7^ EVs, equivalent to a concentration of 5x10^8^ EVs per mL), also at different doses, were administered. NO production was measured, and a reduction in NO levels reflected an anti-inflammatory effect in a dose-dependent manner.

Collectively, these results illustrate that applying hydrodynamic stimulation to producer cells within our system not only enhances EV production and increases yield but also results in EVs with superior functional properties in wound healing, neo-angiogenesis, and anti-inflammatory activity. Furthermore, the cellular 3D organization emerges as a crucial factor influencing these activities. EVs derived from spheroids (HydroSph and Starv 3D) exhibit enhanced functional efficacy compared to those from 2D-cultured cells (HydroCell and Starv 2D), underscoring the importance of utilizing cells in a 3D configuration. This approach not only improves the functional attributes of the EVs but also better replicates the in vivo environment, thereby providing a more physiologically relevant context for their application.

### Towards an all-in-one approach: in situ production of spheroids in rotating tubes, followed by EV production

One of the advantages of the technology is its complete modularity: we can design any internal obstacle, then set the rotation speed between 1 and 2000 rpm, the acceleration after each stop and change of direction between 60 and 6000 rpm, the frequency of direction changes (any frequency possible), and the presence of a pause between each change of direction (pause time from null to hours, typically 1-20 s). For EV production, we found that the optimal rotation speed was 600-800 rpm, with a quick direction change (5-second period, meaning 2.5 seconds in each direction) recommended. Our hypothesis was that at slower rotations, with pauses, the tube could be used upstream for spheroid formation, leading to an all-in-one approach of in situ spheroid formation and direct EV production from these spheroids. This approach is illustrated by the schematic in Fig. 4a. For the spheroid formation phase, the rotation speed was set at 60 rpm (1 rotation per second) with an acceleration of 120 rpm, direction change periods ranging from 2 to 40 seconds, and a pause between each direction change ranging from 1 to 20 seconds (including no pause). Starting from individual cells, simply resuspended in the tubes, the resulting spheroids were analyzed through image processing, with diameter measurements. Fig. 4b shows the distribution of obtained diameters, with all diameters below 15 µm corresponding to still individual cells (no spheroid formation), and the corresponding percentage is also indicated in the figure. Images of the obtained spheroids are illustrated in Fig. 4c, with additional images for the same rotation parameters presented in Supplementary fig. 16. The best conditions identified involved short 2-second pauses and rotation periods of 10-20 seconds; in these conditions, almost no cells remained individual (less than 6%), and we avoided overly large structures exceeding 100 µm in diameter. For the subsequent phase, the maturation protocol selected was 10 seconds of rotation in one direction, a 2-second pause, and then 10 seconds of rotation in the reverse direction.

Next, for the EV production phase, the quantification of the number of EVs per producing cell is presented in the graphs of Figures 4d and 4e. These graphs correspond to the same conditions as Figures 1i and 1j, where EVs were produced from spheroids prepared outside the tubes (HydroSph) in microwells. The production rates (Fig. 4d) and spheroid viability (Fig. 4e) are very similar for spheroids prepared in situ in the tubes (HydroSphIn), with optimal production in terms of yield versus viability around 600 rpm. The advantage of having the spheroids in situ in the tube is the ability to alternate between production phases and rest/maturation phases for the spheroids. This is shown in Fig. 4e (production) and Fig. 4f (viability), where 2-hour production periods alternate with rest periods (spheroid maturation protocol) of 6 to 12 hours. At 600 rpm, the highest production rates are achieved, with 15,000 EVs per producing cell (as measured by Videodrop, equivalent to more than 60,000 EVs per cell by the standard NanoSight measurement), a rate rarely, if ever, matched by other production methods. However, mortality increases after the second stimulation but remains below 40%, while it can reach up to 80% for 1600 rpm stimulations, which are also lower in terms of production. Conditions at 200 rpm and 400 rpm are interesting, especially for more fragile cells, with significant production rates exceeding 5,000 EVs per cell and very reasonable mortality.

Clearly, the production of EVs from spheroids formed in situ presents a significant advantage in the hydrodynamic bioproduction technology. However, it was still necessary to evaluate the impact of the formation mode on spheroid biology. To this end, we analyzed the gene expression profiles of hMSC spheroids formed either in microwells or directly in situ. These analyses were conducted both after spheroid formation (before stimulation) and following the stimulations, either hydrodynamic or static starvation. Various mesenchymal, epithelial, and anti-apoptotic markers were assessed using RT-qPCR. Overall, these markers were present in all spheroid conditions and were consistently over-expressed compared to the 2D monolayer control, indicating a retained mesenchymal signature regardless of formation mode or stimulation applied. Comparing spheroids formed in microwells to those formed directly in situ revealed no significant differences, suggesting that the formation process does not impact spheroid maturation; in situ-formed spheroids are comparable to those formed externally.

Hydrodynamic stimulation slightly increased the expression of most mesenchymal markers, as well as the anti-apoptotic marker MCL1. This increase was also observed post-starvation, albeit to a higher degree, especially for the MCL1 marker. These findings suggest that stimulations do not drastically alter the genomic signature of the spheroids but may have a slight impact on their biology. Notably, the impact of hydrodynamic stimulation appears less significant compared to starvation, indicating that hydrodynamic stimulation better preserves the biological integrity of the spheroids than traditional starvation methods used for EV production.

**Fig. 4:**
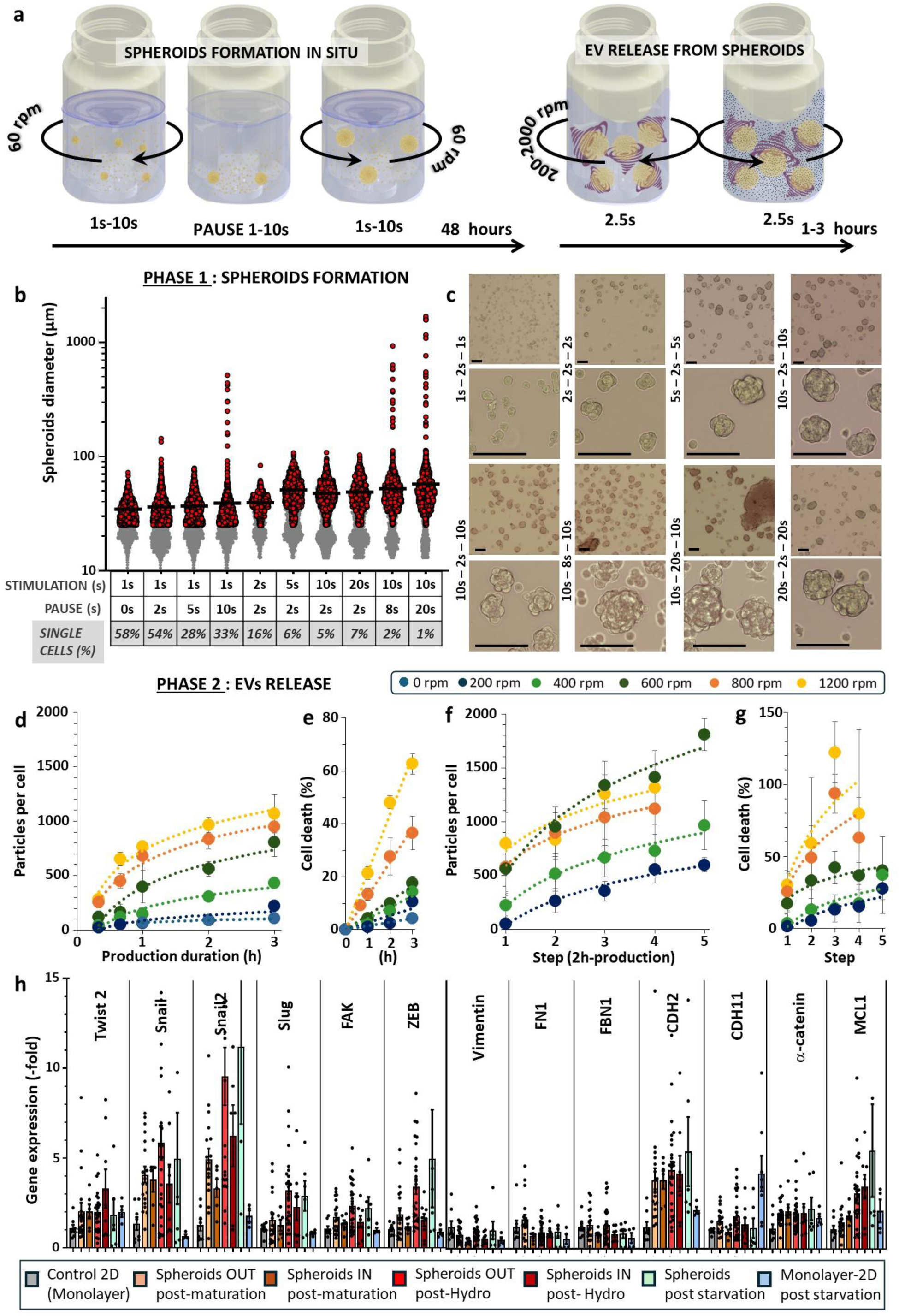
Streamlined bioproduction: Spheroid formation and mass EV release via tuning of stimulation parameters. **a**: Schematic diagram illustrating the all-in-one production approach. The process begins with single cells, progresses through their aggregation into spheroids, and culminates in the release of EVs. **b**: Diameter distribution of the produced spheroids (at least 200 spheroids measured for each condition), achieved using specific configurations of rotation per minute (rpm) and duration for each step, as displayed in the accompanying chart. **c**: Representative images of spheroids produced under each condition, corresponding to the configurations shown in b. **d**: Quantification of EV production, presented as the number of particles produced per producing cell, under different stimulation conditions (varied rpm) and durations (up to 3 hours). **e**: Induced cell mortality under the same conditions as (D). **f**: EV production with a stepwise protocol of 2-hour production periods interspersed with 6 to 12 hours slow rotation, showing the number of particles produced per cell for different stimulations. **g**: Cell mortality associated with the conditions described in (F). **h:** Expression of a panel of mesenchymal and apoptotic genes measured by RT-qPCR for control cells cultured in monolayers (Control 2D), after the formation of spheroids either in microwells (Spheroids Out post-maturation) or in baffled tubes (Spheroids In post-maturation), and after stimulations: hydrodynamic for spheroids Out and In (Spheroids Out post-hydro & Spheroids In post-hydro), and starvation in monolayers (Starv2D).

### EVs produced from spheroids formed in situ maintain excellent functionality

The standout feature of this system is its capacity to streamline both spheroid production and their large-scale release of EVs. Consequently, it was crucial to evaluate the biological properties and therapeutic potential of these EVs produced by spheroids formed in situ (SphIn). The same analyses presented in Fig. 1 are shown in Fig. 5 for the condition HydroSphIn, revealing structurally intact EVs with similar characteristics, as observed by cryo-TEM (Fig.5a, along with additional images in Supplementary fig.9). The expression of CD63, CD81, and Syntenin-1 (SYNT-1) was confirmed via Western blot (Supplementary fig. 11C), while the expression of CD63, CD81, CD29, CD44, CD49e, CD105, and CD146 was verified by MACSplex analysis (Supplementary fig. 12). For all analyses, similar expression levels were observed for the HydroSphIn condition compared to all other conditions, particularly showing comparable marker expression to the HydroSph condition. For the functional assays, Fig. 5b displays a representative image of wound closure induced by SphIn-derived EVs, showing significant efficacy (with additional images in Supplementary fig. 13I), and Fig. 5c quantifies the healing kinetics, revealing comparable effectiveness to 10% FBS, and statistically superior healing at 72 hours. Fig. 5d demonstrates sprouting from endothelial cell spheroids in response to SphIn-derived EVs, with the effect quantified in Fig. 5e, showing superior efficacy to VEGF. Lastly, Fig.5f highlights the robust anti-inflammatory activity of EVs produced from the spheroids formed in situ under hydrodynamic conditions.

**Fig. 5:**
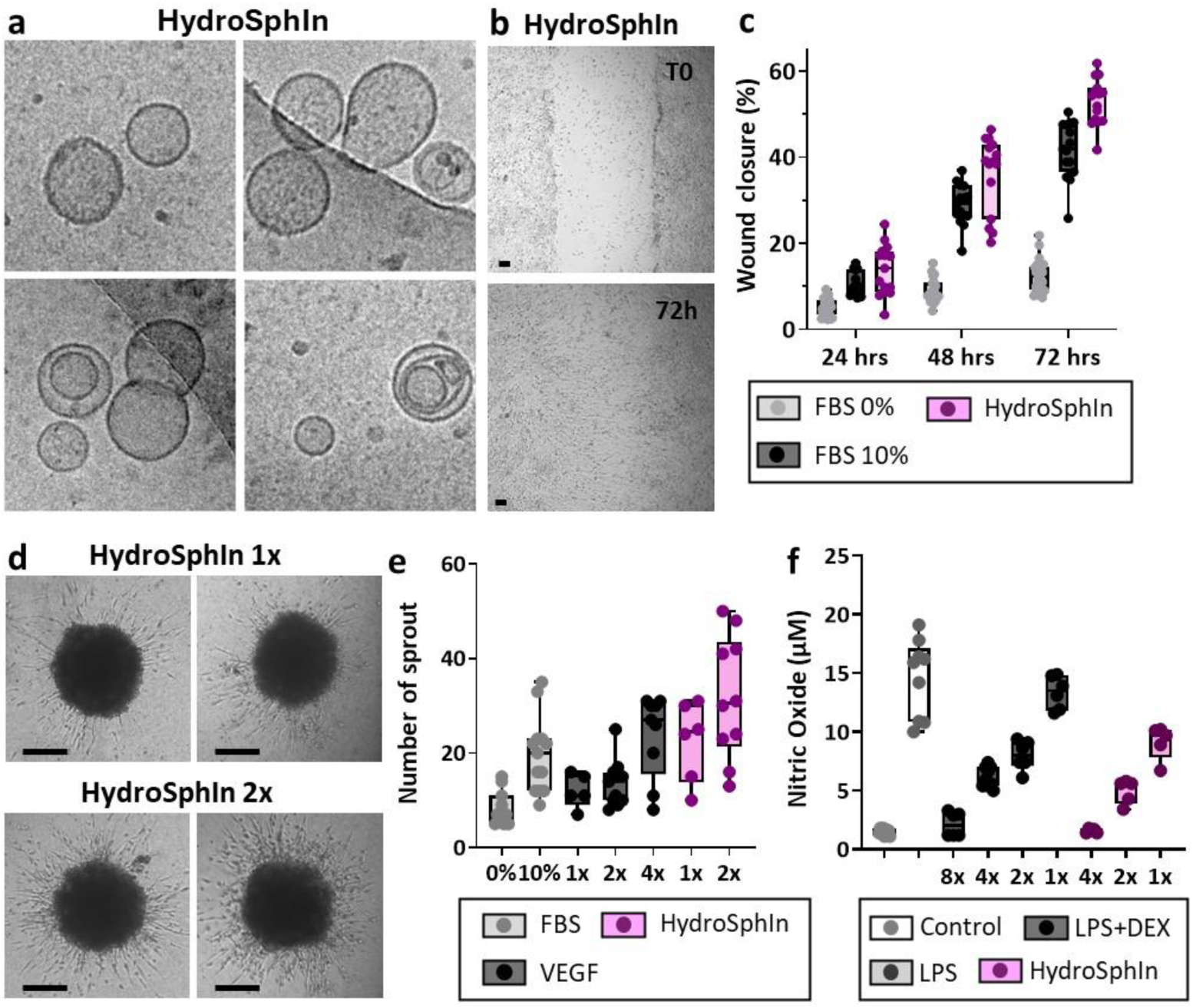
Characterization of EVs produced hydrodynamically from in-tube formed spheroids (HydroSphIn). **(A)** Morphology of EVs observed by CryoTEM. **(B)** MacsPlex quantification, similar to Figures 2F, with blue and green points representing same control conditions (Starv2D and Starv3D), and purple points indicating independent HydroSphIn production conditions. **(C)** Representative wound healing images for a HydroSphIn condition at the initial wound and after 72 hours. Scale bar = 100 µm. **(D)** Quantification of wound healing effect, similar to the graph presented in Fig.4B. Controls (FBS 0% and 10%) are the same, with purple points indicating independent measurements for HydroSphIn. **(E)** Images of neo-angiogenesis for two doses of HydroSphIn EVs. Scale bar = 100 µm. **(F)** Quantification of the number of sprouts, with the same FBS and VEGF controls as in Fig.4D. Purple points represent independent measurements for HydroSphIn. **(G)** Anti-inflammatory effect of HydroSphIn EVs (independent purple points), with the same FBS, LPS, and LPS+DEX controls as in Fig.4E.

### Hydrodynamic stimulation of stem cell spheroids: game changer in EV manufacturing?

The need for reliable and scalable methods for extracellular vesicle (EV) production, isolation, characterization remains critical due to the ever-increasing advancements in EV research within the scientific and biotechnological fields^74^. In response, our study introduces a novel approach that leverages a hydrodynamic stress to trap spheroids and stimulate them to release EVs at high rate. This innovation provides a new framework for controlling and enhancing the bioproduction process of EVs through hydrodynamic stimulation, overcoming a long-standing challenge of improving production yields. The ability to finely adjust key production parameters addresses the persistent need for more efficient, scalable, and robust EV manufacturing processes.

This breakthrough hydrodynamic approach not only increases the quantity of high-quality EVs but also reveals a unique proteomic profile for EVs generated under flow conditions. From a pharmaceutical manufacturing standpoint, the hydrodynamic flow parameters introduced can thus be regarded as critical process parameters (CPP). These variables significantly influence the production process, and therefore must be monitored or controlled to ensure the quality of the final product, as specified by regulatory frameworks such as ICH Q8 (R2). The relationship between process parameters and the critical quality attributes (CQA) of the product is essential for optimizing production, and we believe this methodology will facilitate the development of more precise design spaces in pharmaceutical manufacturing.

The journey toward regulatory approval and the commercialization of EV-based therapies faces numerous challenges, particularly in establishing efficient, reproducible, and controllable production systems. While academic research continues to push the boundaries of EV science, the lack of robust and scalable production methods remains a significant barrier. This ongoing work aims to address this gap by offering a method that can bridge the academic and industrial divides and meet the demands of regulatory agencies.

At the current stage of development, the technology’s ability to streamline spheroid production and EV release, coupled with the specific proteome signature and enhanced activity of the EVs produced, paves the way for scaling up EV production for therapeutic applications.

## CONCLUSION

The extracellular vesicles (EVs) generated through hydrodynamic stimulation in the reverse rotating baffled tubes, from stem cells and spheroids, retain the essential characteristics of EVs, including their morphology, size, and proteins classified as MISEV classes 1 and 2. Additionally, they exhibit a unique cargo. This cargo is influenced not only by the hydrodynamic stimulation but also by the specific physiological environment of stem cells organized as tissue-like spheroids. Proteomic analysis revealed an upregulation of proteins associated with angiogenesis and wound healing within these EVs. This upregulation translated into functional benefits, such as improved sprout formation from endothelial cells and accelerated wound closure with dermal fibroblasts. Furthermore, proteins associated with mitochondria were found to be significantly upregulated under hydrodynamic conditions. This overexpression is likely responsible for the enhanced anti-inflammatory activity observed in the hydrodynamic EVs. Clearly, organizing stem cells into spheroids within a tissue-like environment closer to their natural setting in the body provides additional functionalization. However, pre-organizing the cells into spheroids using microfabrication techniques, while feasible, adds an extra, time-consuming, and relatively costly step, which could hinder scalability. Remarkably, the reverse rotating tube technology also allows for the formation of spheroids in the first step within the tube itself, thanks to the modularity of the stimulation conditions. These spheroids exhibit the same mesenchymal genetic profile as those formed outside the tubes, a profile that remains largely unchanged after hydrodynamic stimulation, regardless of the spheroids used. The EVs produced by these in situ spheroids are also very similar to those produced by their microfabricated counterparts, both in terms of characterization and function. This dual applicability of spheroid formation and EV release within the same technology certainly makes it a competitive solution for enabling tailored EV production processes to meet specific therapeutic requirements, offering high-yield manufacturing and scalability.

## ACKNOWLEDGEMENTS

This work was supported by the European Union (ERC-2019-CoG project NanoBioMade 865629), and Institut Pierre-Gilles de Gennes (Investissements d’avenir ANR-10-EQPX-34). We thank Jean-Michel Guignier (Sorbonne Université, IMPMC, Paris) for CryoTEM observations, Nicolas Ansart (CurieCoretech EV, Institut Curie, Paris) for performing WB and MacsPlexExo analyses of EVs, and Patrick Poullet from the Curie bioinformatics platform U900 for the development of myProMS.

## METHODS

### Cell Lines and Cell Culture

The study utilized hTERT-immortalized human Mesenchymal Stem Cells (hMSC; abm, T0523), Human Umbilical Vein Endothelial Cells (HUVEC; ATCC, CRL1730), and hTERT-immortalized Human Skin Fibroblasts (HSF; abm, T0326). All cell lines were cultured in a humidified atmosphere of 5% CO_2_ at 37°C (incubator MCO-201IC, Panasonic), and all culture media were supplemented with 1% penicillin-streptomycin (Gibco, 15140-122). Specifically:

hMSCs: Cultured in T150 flasks (TPP, 90150) with culture medium Prigrow II (abm, TM002) supplemented with 10% Fetal Bovine Serum (FBS) (Gibco, 12662029), and 1µM hydrocortisone (Sigma, H0135).

HUVECs: Cultured in T75 flasks (TPP, 90075) in DMEM (Gibco, 21885) supplemented with 10% heat-inactivated FBS (Dutscher, S1900-500C).

HSFs: Cultured in PriCoat T25 flasks (abm, G299) in culture medium Prigrow III (abm, TM003) supplemented with 10% FBS (Gibco, A5209402).

### Spheroids formation in microwells

Spheroids were formed in microwells using custom 3D-printed circular molds. Each mold featured 2269 pillars, each 200 µm in height and diameter, designed to fit into wells of a 6-well plate. These molds were created with a DigitalWax 028J Plus 3D printer and DigitalWax DS3000 resin. To prepare the microwells, a solution of 2% agarose (Sigma, A0576) in PBS was heated and poured into the wells of a 6-well plate. The 3D-printed molds were carefully placed in contact with the agarose solution, and the assembly was cooled at 4°C for 20 minutes to allow the agarose to solidify. After gelation, the molds were removed, and the plates were sterilized under UV light for 1 hour and stored at 4°C in PBS until use. For spheroid formation, hMSCs (passages 10-20) were detached from their culture flasks, centrifuged at 300g for 5 minutes, and resuspended at a concentration of 700,000 cells/mL in complete medium. 1 mL of this cell suspension was added to each well containing the agarose microwells. The plates were incubated at 37°C for 30 minutes to allow cells to sediment into the microwells, followed by the addition of 2 mL of complete medium per well. The cells were then incubated at 37°C for 24 hours to form cohesive spheroids.

Prior to use, spheroids were recovered from the microwells by gentle pipetting, followed by three washing steps. Each wash involved centrifugation at 200 g for 3 minutes and resuspension in 220-nm filtered DMEM without phenol red (Gibco, 31053-028). The final suspension was then ready for EV production.

### EVs production under starvation conditions from monolayers and spheroids

*From Monolayers*: To produce EVs from monolayers (starvation 2D), cells were grown in T150 flasks (TPP, 90150) until they reached confluency (approximately 10 million cells per flask, equivalent to 500,000 cells per mL). The culture medium was then replaced with 20 mL of fresh serum-free medium, and the flasks were incubated at 37°C with 5% CO2 for 72 hours. After this incubation period, the conditioned medium containing EVs was collected and centrifuged at 2000 g for 10 minutes at 4°C for further processing.

*From Spheroids*: For the production of EVs from spheroids (starvation 3D), T150 flasks were pre-coated with 20 g/L agarose diluted in PBS to prevent cell adhesion. A suspension of spheroids was prepared as previously described, with a density ranging between 500 and 2000 spheroids per mL (approximately 150,000 to 600,000 cells per mL). Each flask was filled with 20 mL of this spheroid suspension and incubated at 37°C with 5% CO2 for 72 hours. Following the incubation, the medium containing spheroids and EVs was collected and centrifuged at 300 g for 5 minutes to remove the spheroids. The resulting conditioned medium was then further centrifuged at 2000 g for 10 minutes at 4°C for subsequent use.

### Reverse rotating baffled tube device

A prototype of the device was developed to optimize an all-in-one approach for the technology. This initial prototype incorporates 3D-printed tubes, motors, electronics, an incubator for thermal and CO2 regulation, and a parameter control program. A photograph of the prototype is available in Supplementary fig.17.

*3D printing of the tubes*: The tubes were printed using a DigitalWax 028J Plus 3D printer with LOCTITE 3D MED412 resin, which meets ISO 10993-5 and -10 standards for biocompatibility.

*Motors*: Stepper motors (RS PRO, 2.8 V, 5mm shaft diameter) were used for this prototype. These motors are easily controllable, robust, and cost-effective. They were mounted on an aluminum breadboard (250 mm x 250 mm, Thorlabs), with six motors per board.

*Electronics and Programming*: The motor microcontrollers were attached to PCBs to optimize space and wiring. A Raspberry Pi microcomputer was used to generate the control interface, and the interface software was coded in Rust.

*Incubator*: One disadvantage of the stepper motors used in this prototype is the heat they generate when powered. To ensure proper regulation of various parameters (temperature and gas), the aluminum breadboard with the motors was placed inside a small Peltier-cooled CO2 incubator (WCI-40P model, INSTRUMAT), with chamber dimensions of 320x350x375mm.

### Spheroids formation and hydrodynamic EV production in reverse rotating baffled tubes

This novel device for extracellular vesicle (EV) production boasts complete modularity: it allows for adjustable rotation speeds (1–6000 rpm), inversion frequencies (0.01–10 Hz), and optional pauses (1–60 seconds). These parameters govern the establishment of internal hydrodynamic flows, enabling an all-in-one approach for different regimes, from cell suspension cultures and in situ spheroid formation to EV production from cells and spheroids.

*Spheroid Formation in Tubes:* Cells are suspended in their complete culture medium at a density ranging from 200,000 to 500,000 cells per mL. The stimulation parameters to be tested vary between 40 and 80 rpm for rotation speed, 0.05 to 1 Hz for inversion frequency, and 0 to 20 seconds for pauses between rotations. Under certain conditions, cells are cultured in suspension without forming spheroids.

*EV Production in Tubes:* For EV production, the producer cells, either as individual cells or spheroids (formed in microwells or directly in the tubes), are harvested and subjected to three washes with serum-free DMEM to remove any serum residues that could interfere with particle measurement. Cells are then resuspended in 20-25 mL per tube at densities equivalent to 200,000–500,000 cells per mL (approximately 500–2000 spheroids per mL). The reverse rotation protocol is initiated (200-1600 rpm, 0.2 Hz, no pause), typically for a maximum of 3 hours, with optional pauses (up to 20 seconds each) to collect 100 µL samples for particle measurement. Sequential stimulation is also possible, alternating rapid phases (200–1200 rpm, 0.2 Hz) for 2 hours with maturation phases (60 rpm, 0.05 Hz, 2-second pauses) for 6-12 hours. At the end of the production process, the tube volume is collected, spheroids or cells are removed by centrifugation at 300 g for 5 minutes, and the conditioned medium is prepared by centrifugation at 2000 g for 10 minutes at 4°C for further processing.

### Viability assays

*Live/Dead imaging of spheroids*: 100 µL of Live/Dead reagent (Invitrogen 488/570) were added to 50 µL of spheroids suspension (containing in between 100 and 1000 spheroids) in 96-well plate (PerkinElmer, Optiplate-96) and left to incubate for 30 min at room temperature. Spheroids were imaged by plate reader (Ensight Multimode Plate Reader, PerkinElmer) in bright field, and in fluorescence (excitation 465 nm and 535 for Live (green) and Dead (orange) respectively. Fiji was used to quantify the fluorescence levels on region of interest delimited by the spheroids outline.

*Toxilight cell death assay*: 20 µL of Toxilight reagent (Lonza) were combined with 5 µL of the conditioned media in 384-well plate (PerkinElmer, Optiplate-384) and left for 15 min at room temperature. Luminescence was next measured by plate reader (0.1 measurement time). Positive control was obtained by lysis of spheroids or cells at same concentration.

*AlamarBlue metabolic activity assay*: Cells or spheroids were placed into a 96-well plate, with a density of approximately 200-800 spheroids per well or 5000-20000 cells per well. Control samples were seeded at the same density for each experiment. Each well was then incubated with 200 µL of phenol red-free DMEM supplemented with 10% Alamar Blue (Invitrogen) for 2 hours. After incubation, 100 µL of the medium from each well was transferred to a new 96-well plate. Fluorescence was measured using Ensight plate reader, with an excitation wavelength of 570 nm and an emission detection wavelength of 585 nm.

### Nanoparticles Tracking Analysis (NTA): NanoSight measurements

NTA was performed on conditioned medium samples using the NanoSight NS300 (Malvern) with a 532 nm laser module. Prior to analysis, if needed, samples were diluted to achieve a maximal concentration of 2x10^8^ particles/mL. For each measurement, 1 mL of the diluted sample was injected into the apparatus chamber at a flow rate of 20-30 μL/min using a sterile 1-mL syringe. Baseline measurements, including buffer and time zero before production, were systematically performed. Each sample underwent five 1-minute acquisitions, captured with the camera level set to 16. The recorded videos were subsequently analyzed using NanoSight NTA software, maintaining a detection threshold of 4.

### Nanoparticles Tracking Analysis: Videodrop measurements

Interferometric Light Microscopy (ILM) using the Videodrop instrument (Myriade, Paris) was employed to analyze nanoparticle concentration and hydrodynamic diameter by recording the diffraction patterns created by nanoparticles moving in the light path. Prior to each measurement, the sample chip was thoroughly cleaned with ethanol and distilled water.

For the measurements, an 8-μL sample was placed at the center of the sample chip. The chip was then positioned in the optical path of the Videodrop instrument. Samples were analyzed directly at their concentration in conditioned medium, with an effective working range of 10^8^-10^10^ particles per mL. At least duplicate measurements per samples were performed. The ILM signal enabled nanoparticle detection and tracking, with the data processed using the qvir software, with the doublet detector for all measurements. Summary reports were exported as PDFs, and raw data were saved in .qvir and .csv formats. Measurements were also conducted at time zero, before production, to serve as a baseline.

### EV isolation by Size Exclusion Chromatography (SEC)

To concentrate the conditioned media, 100 kDa cut-off centrifugal filters (Sigma, Centricon Plus-70 100kDa, UFC710008) were utilized, yielding 300-500 µL of concentrated conditioned media (CCM), in accordance with the manufacturer’s instructions. EVs were subsequently isolated and purified using size exclusion chromatography (SEC). For this process, SEC columns (Izon Sciences, qEVoriginal/70nm Gen2 SEC columns, ICO-70) were pre-washed with 25.5 mL of PBS. Each sample’s 500 µL of CCM was then carefully applied to the top of the columns. After the entire sample entered the column, PBS was added. The initial 2 mL fraction was discarded, and the following 2.4 mL fraction, containing the EVs, was collected. In certain cases, a subsequent 2.4 mL fraction (Intermediate fraction) was also collected. Both EV and Intermediate fractions were then reconcentrated using 10 kDa cut-off centrifugal filters (Milipore, Amicon Ultra-4, UFC8010), resulting in 100 µL of concentrated EV suspension (EV pool). The EV pools were either stored at 4°C for immediate analysis or transferred at -80°C for long-term storage.

### Cryogenic Transmission Electron Microscopy (cryoTEM) imaging

A 5 μL sample aliquot was applied to a quantifoil carbon membrane grid (Quantifoil Micro Tools GmbH). Excess liquid was blotted from the grid for a few seconds, followed by rapid plunging into liquid ethane to form a thin vitreous ice film. The grid was then mounted onto a Gatan 626 cryo-holder, cooled with liquid nitrogen, and inserted into a JEOL JEM2100 (LaB6) electron microscope operating at 200 kV. The temperature was maintained at -180 °C throughout the imaging process. Images were captured using an Ultrascan 1000 CCD camera (Gatan) at a resolution of 2k x 2k pixels.

### Protein quantification

The protein content of EVs was quantified utilizing the Micro BCA Protein Assay Kit (Thermo Fisher Scientific). To establish a standard curve, Bovine Serum Albumin was diluted to final concentrations of 0.5, 1, 2.5, 5, 10, 40, and 200 µg/mL. In parallel, EVs were mixed with RIPA buffer (Thermo Fisher Scientific) at a 1:7 dilution and incubated on ice for 30 min for EV lysis. EVs lysates were then diluted in phosphate buffer saline (PBS) to reach a final 1:10 RIPA concentration. 45 µL of each EVs sample and each standard were then transferred in triplicates into a 384-well microplate (PerkinElmer, Optiplate-384) and 45 µL of the BCA working reagent was added to each well. The plate was then agitated on a plate shaker for 30 seconds, covered, and incubated at 37°C for 2 hours. Absorbance readings at 562 nm were taken using a Ensight plate reader, and protein concentrations were calculated based on the standard curve.

### Whole proteome analysis

Five biological replicates of each analyzed condition were assessed by Liquid Chromatography coupled with Tandem Mass Spectrometry (LC-MS/MS) using 5µg of proteins of each replicate.

*Sample Preparation for LC-MS/MS*: Protein pellets of 5 µg each were dried under vacuum using a Savant Centrifuge SpeedVac concentrator (Thermo Fisher Scientific). The dried pellets were then solubilized and reduced in 10 µL of 8 M urea, 100 mM ammonium bicarbonate, and 5 mM dithiothreitol (DTT) at pH 8.0, and incubated at 57°C for 1 hour. Following this, the samples were allowed to cool to room temperature, after which iodoacetamide was added to a final concentration of 10 mM. The alkylation reaction was conducted in the dark for 30 minutes at room temperature. The samples were subsequently diluted to a final urea concentration of 1 M using 100 mM ammonium bicarbonate at pH 8.0. Protein digestion was performed using trypsin/LysC (1 µg, Promega) in a total volume of 100 µL, with the mixture incubated overnight at 37°C under constant vortexing. To desalt the digested peptides, the samples were passed through homemade C18 StageTips. Elution of peptides was achieved using a solution of 40% acetonitrile (CH3CN) and 60% water containing 0.1% formic acid. The eluate was then concentrated to dryness under vacuum.

*LC-MS/MS Analysis*: The peptide samples were analyzed using a Vanquish Neo LC system (Thermo Scientific) connected to an Orbitrap Astral mass spectrometer, utilizing a Nanospray Flex ion source (Thermo Scientific). Separation was achieved on a C18 column (75 µm inner diameter × 50 cm, 2 µm particle size, 100 Å pore size, Thermo Scientific) maintained at 50°C. Peptides were eluted using a linear gradient of 100% buffer A (0.1% formic acid in water) to 28% buffer B (0.1% formic acid in acetonitrile) over 104 minutes at a flow rate of 300 nL/min. Ionization was performed with a spray voltage of 2200V, funnel RF level set to 40, and a heated capillary at 285°C. Full MS scans were acquired in the range of 380–980 m/z with a resolution of 240,000 at m/z 200, using a normalized AGC target of 500% and a maximum injection time of 5 ms. Fragmentation was carried out in data-independent acquisition (DIA) mode, covering a precursor mass range of 380–980 m/z with 2 Da isolation windows. Fragment ions were generated in the HCD cell with a normalized collision energy of 25%, a normalized AGC target of 500%, and a maximum injection time of 3 ms.

*Data Processing LC-MS/MS*: Data analysis was conducted using the Pulsar search engine integrated with Spectronaut v19 (Biognosys) to perform directDIA+ analysis on the collected LC-MS/MS data. The raw data were searched against the Homo sapiens (UP000005640) Uniprot database. The search parameters included trypsin enzyme specificity, allowing up to two missed cleavage sites. Carbamidomethylation of cysteine was specified as a fixed modification, while N-terminal acetylation and methionine oxidation were considered variable modifications. The processed data were further analyzed using myProMS v3.10 (available at https://github.com/bioinfo-pf-curie/myproms)^75^. Protein quantification was performed by extracting ion chromatograms (XICs) of proteotypic peptides common to the conditions being compared (TopN matching). The analysis permitted missed cleavages and included carbamidomethylation. Peptide-level median and scale normalization was applied to adjust the total signal across the five biological replicates. To assess the significance of protein abundance changes, a linear model adjusted for peptides and biological replicates was employed. A two-sided T-test was conducted on the fold change estimates derived from the model, and p-values were corrected using the Benjamini-Hochberg FDR method. Label-Free Quantification (LFQ) was also performed according to a specified algorithm^76^, requiring a minimum of two peptide ratios and utilizing the large ratios stabilization feature. Proteins were considered significant if they were identified with at least two distinct peptides across five biological replicates and exhibited a log2(fold change) ≥ 2 or ≤ -2 with an adjusted p-value < 0.05. Proteins with a log2(fold change) between -2 and 2 were deemed common between the conditions. The raw mass spectrometry proteomics data can be found in the ProteomeXchange Consortium via the PRIDE partner repository^77^.

### Transcriptomic analysis with RT-qPCR

hMSCs cells or spheroids were collected either post-maturation (after spheroids formation and before stimulation), post-rotation (after stimulation) or post-starvation (no rotation stimulation). Total RNA of those samples was extracted using NucleoSpin RNA mini kit (Machery-Nagel, Thermo Fisher Scientific, #740955.50) following the manufacturer instructions. To avoid contaminations with genomic DNA, RNA samples were incubated with desoxyribonuclease (DNAse) allowing this unwanted genomic DNA degradation. The extracted RNA concentrations and purities were then assessed using the Nanodrop device (Ozyme).

Reverse transcription was performed from 1µg of the previously extracted RNA using the High-Capacity cDNA Reverse Transcription kit (Thermo Fisher Scientific, #4368814) with random primers, according to the manufacturer protocol. The obtained cDNA was then quantified using the same Nanodrop device.

Real-time quantitative PCR was carried out, starting with 30ng of cDNA using Power SYBRGreen qPCR Master Mix (Thermo Fisher Scientific, #A25742), on the QuantStudio 3 instrument (Applied Biosystems). Expression of the acide ribosomal protein P0 (RPLP0) was used as reference gene to normalize the obtained Ct values of the target genes. Next, the normalized Ct values were compared to the control 2D monolayer and gene expression folds were calculated based on the comparative Ct method : 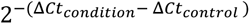.

Mesenchymal markers genes (Twist2, Snail1&2, Slug, FAK, ZEB, Vimentin, FN1, FBN1, CDH2, CDH11), epithelial marker gene (α-catenin) and anti-apoptotic-associated gene (MCL1) were assessed by RT-qPCR. The primer sequences are listed in Table 1.

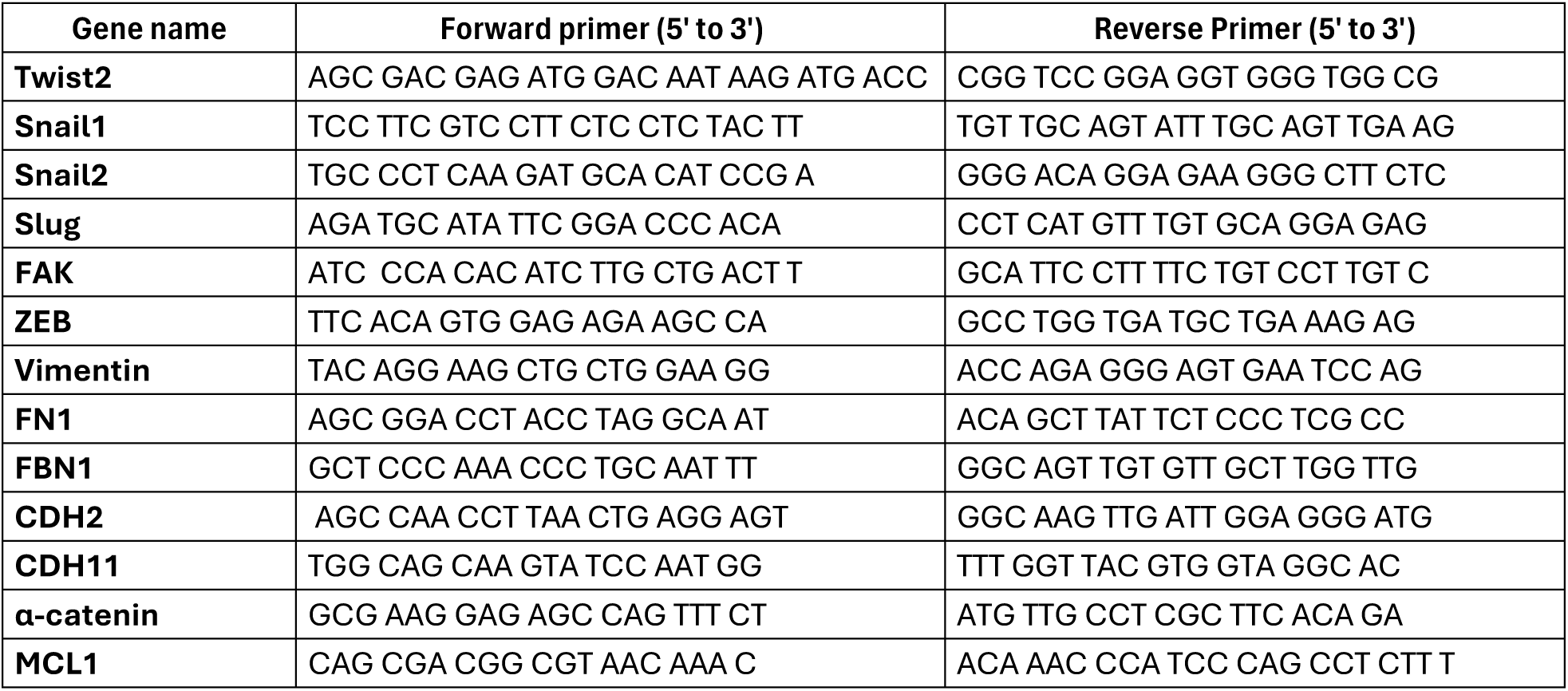

### Wound healing assay

Human skin fibroblasts (HSFs) were seeded into 96-well plates (TPP, 92096) at a density of 100,000 cells per well and cultured at 37°C with 5% CO2 until they reached confluence. A physical wound was created within the monolayer using a 3D-printed template and a sterile 200 µL pipette tip. The culture medium was then replaced with 100 µL of serum-free medium, or serum-free medium supplemented with 1%, 2%, 4%, or 10% fetal bovine serum (FBS), as well as serum-free medium containing extracellular vesicles (EVs) produced under various conditions. The initial EV concentration was set at 10^9^ EVs/mL. HSFs were incubated for 96 hours at 37°C, 5% CO^2^, with images taken periodically using a PerkinElmer Ensight Multimode Plate Reader. At each time point, the area of cell coverage within the wound gap was measured using Fiji software (see Supplementary fig. 18 for an illustration of the thresholding process). The wound healing progress was quantified by calculating the area covered by cells relative to the total area of the wound.

### Angiogenesis sprouting assay on collagen matrix

HUVEC spheroids were generated using a protocol similar to that for hMSC spheroids, with the use of agarose microwells. However, in this instance, the spheroids were produced individually in a 96-well plate, with each well containing a single micropillar mold with a diameter of 500 µm. Thus, a total of 96 microwells were created in each well of a 96-well plate. Following the formation of microwells using 20 mg/mL agarose (200 µL of agarose poured per well) and sterilization, 10,000 HUVECs were added in 20 µL on top of each microwell. The plate was incubated at 37°C for 30 minutes, then centrifuged at 200g for 3 minutes, and 100 µL of medium was added. The cells were incubated for 3 days to allow for spheroid maturation. A 2 mg/mL collagen type I solution (Corning) was prepared on ice, and 30 µL was added to each well of a 384-well plate. After allowing the collagen to polymerize for 30 minutes at 37°C, HUVEC spheroids were carefully transferred onto the collagen matrix, with one spheroid per well. The plates were kept at room temperature for 20 minutes to allow the spheroids to settle. Subsequently, the culture medium was replaced with 50 µL of one of the following treatments: VEGF-α (Sigma) at concentrations of 12.5 (1x), 25 (2x), or 50 (4x) ng/mL; EV pools produced under various conditions and diluted in DMEM, with two doses of 10^9 (1x) EVs and 2x10^9 (2x) EVs per mL; DMEM supplemented with 10% serum; or DMEM alone as a control. The plates were incubated for 24 hours at 37°C and subsequently imaged using the PerkinElmer Ensight Multimode Plate Reader in bright-field mode. Angiogenic sprouting was quantified by measuring the sprout length and counting the number of sprouts per spheroid using Fiji software.

### Anti-inflammatory in vitro assays

To assess the anti-inflammatory activity of EVs, mouse macrophage cells (RAW 264.7) were cultured in DMEM supplemented with 10% FBS (Gibco 10270-106), 1% penicillin-streptomycin (Gibco 15140-122) at 37°C, 5% CO^2^ and kept at low passage (<P20) and low confluency. Stimulation of macrophages with LPS incubation induces the formation of nitric oxide (NO), an indicator of inflammatory reaction. For the assay, cells were seeded in a 96-well plate at a density of 25,000 cells per well in DMEM supplemented with 5% FBS and 1% penicillin-streptomycin. After 48 hours, culture medium was removed and cells were stimulated with 150 µL of DMEM supplemented with 5% FBS and antibiotics (negative control), completed medium supplemented with LPS 0.5 µg/mL only (positive control), with LPS 0.5 µg/mL and EVs (1x = 7.5 x 10^7^ EVs, 2x = 1.5 x 10^8^ EVs, 4x = 3 x 10^8^ EVs) or with LPS 0.5 µg/mL and dexamethasone (Sigma, D4902)(1x = 1µg/mL) (inhibition control) for 24 hours. At the end of the incubation time, an equal volume of a 2% sulphanilamide in 10% phosphoric acid solution and a 0.2% naphtylethylenediamine in water solution were mixed to obtain the Griess’ Reagent. 50 µL of culture medium supernatant was then mixed with 50 µL the Griess’ reagent and incubated 10 min in the dark at room temperature. To measure the NO production, absorbances at 550 nm (measured wavelength) and 620 nm (reference wavelength) were immediately measured using a plate reader EnSight (Perkin Elmer). Sodium Nitrite (NaNO_2_) was used in different concentrations to build a standard curve.

### Statistical analysis

If applicable, statistical differences between groups were assessed using an unpaired, two-tailed Student’s t-test. P-values were denoted as follows: p > 0.05 (not significant NS), p < 0.05 (significant *), p < 0.01 (significant **), and p < 0.001 (significant **).

## Notes

### Competing Interest Statement

The authors have declared no competing interest.

## REFERENCES

1. El Andaloussi, S., Mäger, I., Breakefield, X.O. & Wood, M.J. Extracellular vesicles: biology and emerging therapeutic opportunities. Nature reviews Drug discovery 12, 347–357 (2013).

2. Witwer, K.W. et al. Defining mesenchymal stromal cell (MSC)-derived small extracellular vesicles for therapeutic applications. Journal of extracellular vesicles 8, 1609206 (2019).

3. Elsharkasy, O.M. et al. Extracellular vesicles as drug delivery systems: Why and how? Advanced Drug Delivery Reviews 159, 332–343 (2020).

4. Maldonado, V.V. et al. Clinical utility of mesenchymal stem/stromal cells in regenerative medicine and cellular therapy. Journal of Biological Engineering 17, 44 (2023).

5. Phinney, D.G. & Pittenger, M.F. Concise Review: MSC-Derived Exosomes for Cell-Free Therapy. STEM CELLS 35, 851–858 (2017).

6. Murphy, D.E. et al. Extracellular vesicle-based therapeutics: natural versus engineered targeting and trafficking. Experimental & Molecular Medicine 51, 1–12 (2019).

7. Zhang, B. et al. HucMSC-Exosome Mediated-Wnt4 Signaling Is Required for Cutaneous Wound Healing. STEM CELLS 33, 2158–2168 (2015).

8. Long, Q. et al. Intranasal MSC-derived A1-exosomes ease inflammation, and prevent abnormal neurogenesis and memory dysfunction after status epilepticus. Proceedings of the National Academy of Sciences 114, E3536–E3545 (2017).

9. Chen, P. et al. Targeted delivery of extracellular vesicles in heart injury. Theranostics 11, 2263–2277 (2021).

10. Shi, L. et al. Mesenchymal stem cell-derived extracellular vesicles ameliorate renal interstitial fibrosis via the miR-13474/ADAM17 axis. Scientific Reports 14, 17703 (2024).

11. Miao, L. et al. Extracellular vesicles containing GAS6 protect the liver from ischemia-reperfusion injury by enhancing macrophage efferocytosis via MerTK-ERK-COX2 signaling. Cell Death Discovery 10, 401 (2024).

12. van de Wakker, S.I. et al. Size matters: Functional differences of small extracellular vesicle subpopulations in cardiac repair responses. Journal of extracellular vesicles 13, 12396 (2024).

13. Seltmann, K. et al. Transport of CLCA2 to the nucleus by extracellular vesicles controls keratinocyte survival and migration. Journal of extracellular vesicles 13, e12430 (2024).

14. Oeller, M. et al. Heparin Differentially Regulates the Expression of Specific miRNAs in Mesenchymal Stromal Cells. International journal of molecular sciences 25 (2024).

15. Soler-Botija, C. et al. Mechanisms governing the therapeutic effect of mesenchymal stromal cell-derived extracellular vesicles: A scoping review of preclinical evidence. Biomedicine & Pharmacotherapy 147, 112683 (2022).

16. Gissi, C. et al. Extracellular vesicles from rat-bone-marrow mesenchymal stromal/stem cells improve tendon repair in rat Achilles tendon injury model in dose-dependent manner: A pilot study. PloS one 15, e0229914 (2020).

17. van de Wakker, S.I., Meijers, F.M., Sluijter, J.P.G., Vader, P. & Baker, A. Extracellular Vesicle Heterogeneity and Its Impact for Regenerative Medicine Applications. Pharmacological Reviews 75, 1043–1061 (2023).

18. Seow, K.S. & Ling, A.P.K. Mesenchymal stem cells as future treatment for cardiovascular regeneration and its challenges. Annals of Translational Medicine 12, 73 (2023).

19. S, E.L.A., Mäger, I., Breakefield, X.O. & Wood, M.J. Extracellular vesicles: biology and emerging therapeutic opportunities. Nature reviews. Drug discovery 12, 347–357 (2013).

20. Roefs, M.T. et al. Cardiac progenitor cell-derived extracellular vesicles promote angiogenesis through both associated- and co-isolated proteins. Communications Biology 6, 800 (2023).

21. Al-Sharabi, N. et al. Osteogenic human MSC-derived extracellular vesicles regulate MSC activity and osteogenic differentiation and promote bone regeneration in a rat calvarial defect model. Stem Cell Res Ther 15, 33 (2024).

22. Lin, H. et al. Therapeutic potential of extracellular vesicles from diverse sources in cancer treatment. European Journal of Medical Research 29, 350 (2024).

23. Rädler, J., Gupta, D., Zickler, A. & Andaloussi, S.E. Exploiting the biogenesis of extracellular vesicles for bioengineering and therapeutic cargo loading. Molecular Therapy 31, 1231–1250 (2023).

24. Zheng, W. et al. Identification of scaffold proteins for improved endogenous engineering of extracellular vesicles. Nature Communications 14, 4734 (2023).

25. Zendrini, A. et al. On the surface-to-bulk partition of proteins in extracellular vesicles. Colloids and Surfaces B: Biointerfaces 218, 112728 (2022).

26. Bader, J., Narayanan, H., Arosio, P. & Leroux, J.-C. Improving extracellular vesicles production through a Bayesian optimization-based experimental design. European Journal of Pharmaceutics and Biopharmaceutics 182, 103–114 (2023).

27. Toribio, V. et al. Development of a quantitative method to measure EV uptake. Scientific Reports 9, 10522 (2019).

28. Krivitsky, V. et al. Ultrafast and Controlled Capturing, Loading, and Release of Extracellular Vesicles by a Portable Microstructured Electrochemical Fluidic Device. Advanced materials (Deerfield Beach, Fla.) 35, e2212000 (2023).

29. Gupta, D., Zickler, A.M. & El Andaloussi, S. Dosing extracellular vesicles. Advanced Drug Delivery Reviews 178, 113961 (2021).

30. Poupardin, R. et al. Advances in Extracellular Vesicle Research Over the Past Decade: Source and Isolation Method are Connected with Cargo and Function. Advanced healthcare materials 13, e2303941 (2024).

31. Benayas, B. et al. Proof of concept of using a membrane-sensing peptide for sEVs affinity-based isolation. Frontiers in Bioengineering and Biotechnology 11, 1238898 (2023).

32. Benayas, B., Morales, J., Egea, C., Armisén, P. & Yáñez-Mó, M. Optimization of extracellular vesicle isolation and their separation from lipoproteins by size exclusion chromatography. Journal of Extracellular Biology 2, e100 (2023).

33. Görgens, A. et al. Identification of storage conditions stabilizing extracellular vesicles preparations. Journal of extracellular vesicles 11, e12238 (2022).

34. 34. Zarovni, N. et al. in Extracellular Vesicles: Applications to Regenerative Medicine, Therapeutics and Diagnostics. (eds. W. Chrzanowski, C.T. Lim & S.Y. Kim) 0 (The Royal Society of Chemistry, 2021).

35. Sun, L. et al. Serum deprivation elevates the levels of microvesicles with different size distributions and selectively enriched proteins in human myeloma cells in vitro. Acta pharmacologica Sinica 35, 381–393 (2014).

36. King, H.W., Michael, M.Z. & Gleadle, J.M. Hypoxic enhancement of exosome release by breast cancer cells. BMC Cancer 12, 421 (2012).

37. Piffoux, M. et al. Extracellular vesicles for personalized medicine: The input of physically triggered production, loading and theranostic properties. Adv Drug Deliv Rev 138, 247–258 (2019).

38. Momen-Heravi, F., Bala, S., Kodys, K. & Szabo, G. Exosomes derived from alcohol-treated hepatocytes horizontally transfer liver specific miRNA-122 and sensitize monocytes to LPS. Scientific Reports 5, 9991 (2015).

39. Zhang, W. et al. HIF-1-mediated production of exosomes during hypoxia is protective in renal tubular cells. American journal of physiology. Renal physiology 313, F906–f913 (2017).

40. Muñiz-García, A. et al. Hypoxia-induced HIF1α activation regulates small extracellular vesicle release in human embryonic kidney cells. Scientific Reports 12, 1443 (2022).

41. Li, J. et al. Serum-free culture alters the quantity and protein composition of neuroblastoma-derived extracellular vesicles. Journal of extracellular vesicles 4, 26883 (2015).

42. Otsuka, K., Yamamoto, Y. & Ochiya, T. Uncovering temperature-dependent extracellular vesicle secretion in breast cancer. Journal of extracellular vesicles 10, e12049 (2020).

43. Headland, S.E., Jones, H.R., D’Sa, A.S.V., Perretti, M. & Norling, L.V. Cutting-Edge Analysis of Extracellular Microparticles using ImageStreamX Imaging Flow Cytometry. Scientific Reports 4, 5237 (2014).

44. Böker, K.O. et al. The Impact of the CD9 Tetraspanin on Lentivirus Infectivity and Exosome Secretion. Molecular therapy : the journal of the American Society of Gene Therapy 26, 634–647 (2018).

45. de Almeida Fuzeta, M., et al. Scalable Production of Human Mesenchymal Stromal Cell-Derived Extracellular Vesicles Under Serum-/Xeno-Free Conditions in a Microcarrier-Based Bioreactor Culture System. Front Cell Dev Biol 8, 553444 (2020).

46. Thompson, W. & Papoutsakis, E.T. The role of biomechanical stress in extracellular vesicle formation, composition and activity. Biotechnology Advances 66, 108158 (2023).

47. Kronstadt, S.M. et al. Mesenchymal Stem Cell Culture within Perfusion Bioreactors Incorporating 3D-Printed Scaffolds Enables Improved Extracellular Vesicle Yield with Preserved Bioactivity. Advanced healthcare materials 12, 2300584 (2023).

48. Morrell, A.E. et al. Mechanically induced Ca2+ oscillations in osteocytes release extracellular vesicles and enhance bone formation. Bone Research 6, 6 (2018).

49. Wu, J. et al. Scale-out production of extracellular vesicles derived from natural killer cells via mechanical stimulation in a seesaw-motion bioreactor for cancer therapy. Biofabrication 14 (2022).

50. Chung, J. et al. Fluid Shear Stress Regulates the Landscape of microRNAs in Endothelial Cell-Derived Small Extracellular Vesicles and Modulates the Function of Endothelial Cells. International journal of molecular sciences 23 (2022).

51. Ridger, V.C. et al. Microvesicles in vascular homeostasis and diseases. Position Paper of the European Society of Cardiology (ESC) Working Group on Atherosclerosis and Vascular Biology. Thrombosis and haemostasis 117, 1296–1316 (2017).

52. Vion, A.C. et al. Shear stress regulates endothelial microparticle release. Circulation research 112, 1323–1333 (2013).

53. Vixège, F. et al. Full-volume three-component intraventricular vector flow mapping by triplane color Doppler. Physics in medicine and biology 67 (2022).

54. Jiang, J., Woulfe, D.S. & Papoutsakis, E.T. Shear enhances thrombopoiesis and formation of microparticles that induce megakaryocytic differentiation of stem cells. Blood 124, 2094–2103 (2014).

55. Jo, W. et al. Microfluidic fabrication of cell-derived nanovesicles as endogenous RNA carriers. Lab on a Chip 14, 1261–1269 (2014).

56. Yan, L. & Wu, X. Exosomes produced from 3D cultures of umbilical cord mesenchymal stem cells in a hollow-fiber bioreactor show improved osteochondral regeneration activity. Cell biology and toxicology 36, 165–178 (2020).

57. Jakl, V. et al. A novel approach for large-scale manufacturing of small extracellular vesicles from bone marrow-derived mesenchymal stromal cells using a hollow fiber bioreactor. Frontiers in Bioengineering and Biotechnology 11 (2023).

58. Jalilian, E. et al. Bone marrow mesenchymal stromal cells in a 3D system produce higher concentration of extracellular vesicles (EVs) with increased complexity and enhanced neuronal growth properties. Stem Cell Research & Therapy 13, 425 (2022).

59. Zhang, Y. et al. Systemic administration of cell-free exosomes generated by human bone marrow derived mesenchymal stem cells cultured under 2D and 3D conditions improves functional recovery in rats after traumatic brain injury. Neurochemistry international 111, 69–81 (2017).

60. Thippabhotla, S., Zhong, C. & He, M. 3D cell culture stimulates the secretion of in vivo like extracellular vesicles. Scientific Reports 9, 13012 (2019).

61. Grangier, A. et al. Technological advances towards extracellular vesicles mass production. Advanced Drug Delivery Reviews 176, 113843 (2021).

62. Madel, R.J. et al. Independent human mesenchymal stromal cell-derived extracellular vesicle preparations differentially attenuate symptoms in an advanced murine graft-versus-host disease model. Cytotherapy 25, 821–836 (2023).

63. Cao, J. et al. Three-dimensional culture of MSCs produces exosomes with improved yield and enhanced therapeutic efficacy for cisplatin-induced acute kidney injury. Stem Cell Research & Therapy 11, 206 (2020).

64. Kink, J.A. et al. Large-scale bioreactor production of extracellular vesicles from mesenchymal stromal cells for treatment of acute radiation syndrome. Stem Cell Res Ther 15, 72 (2024).

65. Mendt, M. et al. Generation and testing of clinical-grade exosomes for pancreatic cancer. JCI Insight 3 (2018).

66. Thouvenot, E. et al. High-Yield Bioproduction of Extracellular Vesicles from Stem Cell Spheroids via Millifluidic Vortex Transport. *Advanced materials (Deerfield Beach*, Fla.), e2412498 (2024).

67. Welsh, J.A. et al. Minimal information for studies of extracellular vesicles (MISEV2023): From basic to advanced approaches. Journal of extracellular vesicles 13, e12404 (2024).

68. Nguyen, V.V.T. et al. Inter-laboratory multiplex bead-based surface protein profiling of MSC-derived EV preparations identifies MSC-EV surface marker signatures. Journal of extracellular vesicles 13, e12463 (2024).

69. Hettich, B.F., Ben-Yehuda Greenwald, M., Werner, S. & Leroux, J.-C. Exosomes for Wound Healing: Purification Optimization and Identification of Bioactive Components. Advanced Science 7, 2002596 (2020).

70. Yu, W. et al. HO-1 Is Essential for Tetrahydroxystilbene Glucoside Mediated Mitochondrial Biogenesis and Anti-Inflammation Process in LPS-Treated RAW264.7 Macrophages. Oxidative medicine and cellular longevity 2017, 1818575 (2017).

71. Xia, L. et al. AdMSC-derived exosomes alleviate acute lung injury via transferring mitochondrial component to improve homeostasis of alveolar macrophages. Theranostics 12, 2928–2947 (2022).

72. Hough, K.P. et al. Exosomal transfer of mitochondria from airway myeloid-derived regulatory cells to T cells. Redox biology 18, 54–64 (2018).

73. Zhao, M. et al. Mesenchymal Stem Cell-Derived Extracellular Vesicles Attenuate Mitochondrial Damage and Inflammation by Stabilizing Mitochondrial DNA. ACS nano 15, 1519–1538 (2021).

74. Loria, F. et al. A decision-making tool for navigating extracellular vesicle research and product development. Journal of extracellular vesicles 13, e70021 (2024).

75. Poullet, P., Carpentier, S. & Barillot, E. myProMS, a web server for management and validation of mass spectrometry-based proteomic data. Proteomics 7, 2553–2556 (2007).

76. Cox, J. et al. Accurate proteome-wide label-free quantification by delayed normalization and maximal peptide ratio extraction, termed MaxLFQ. Molecular & cellular proteomics : MCP 13, 2513–2526 (2014).

77. Perez-Riverol, Y. et al. The PRIDE database resources in 2022: a hub for mass spectrometry-based proteomics evidences. Nucleic acids research 50, D543–d552 (2022).

