## Supporting Information for "Harnessing hydrodynamics for high-yield production of extracellular vesicles from stem cells spheroids with specific cargo profiling"

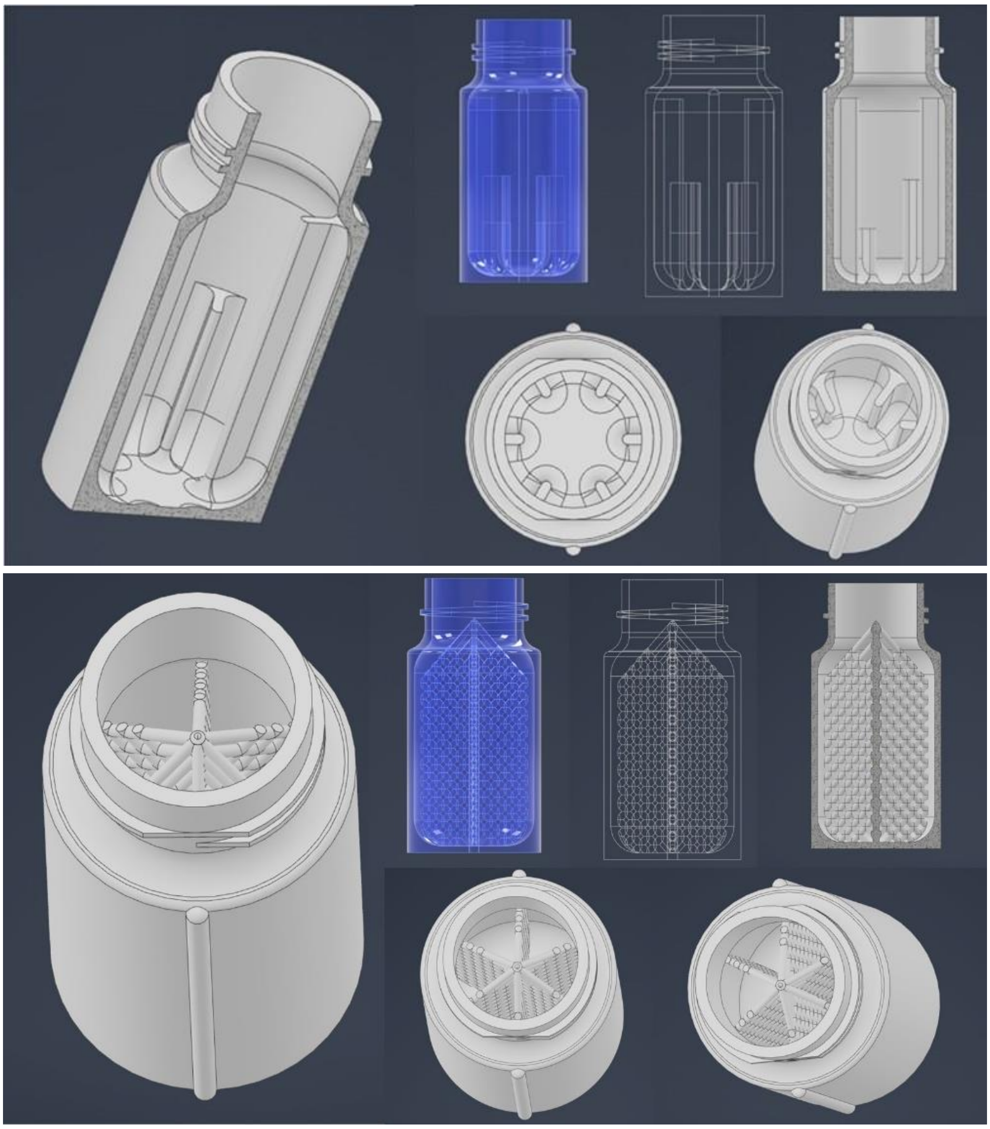

**Supplementary Figure 1** : Concept sketches of the baffled designs, with internal cut walls (A), or internal grids (B) geometries.

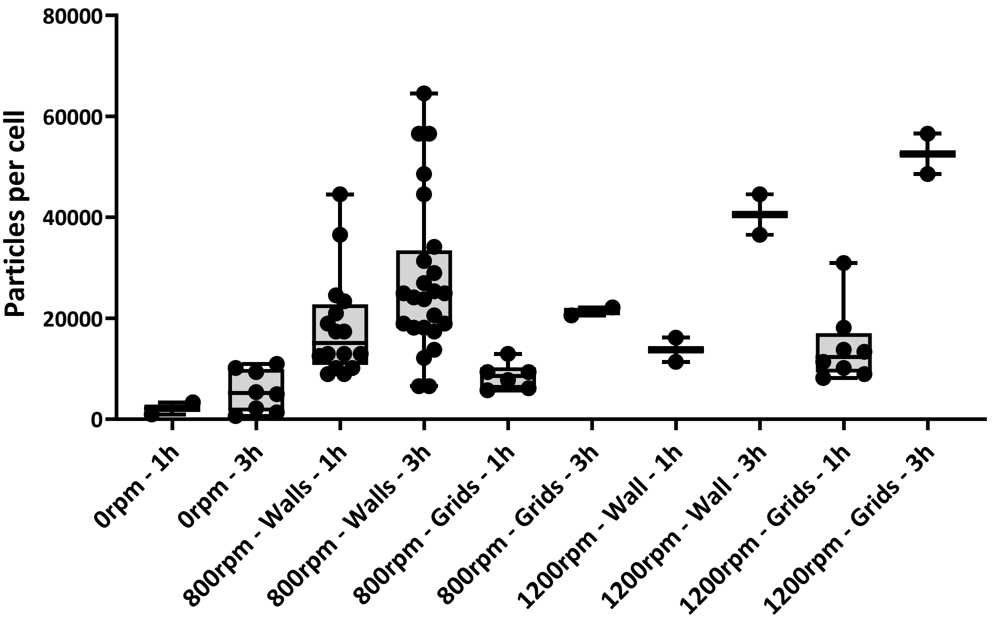


**Supplementary Figure 2**: Comparison of EV production in baffled tubes with internal walls and with grid-like geometries, from spheroids produced in agarose and stimulated at 800 rpm and 1200 rpm. Both tubes geometry have similar efficiency yield, in the range of 30,000 and 50,000 particles per producer cells (forming the spheroids), for 3 hours stimulation at 800 and 1200 rpm, respectively, as measured with NanoSight Nanoparticles Tracking Analysis methods.

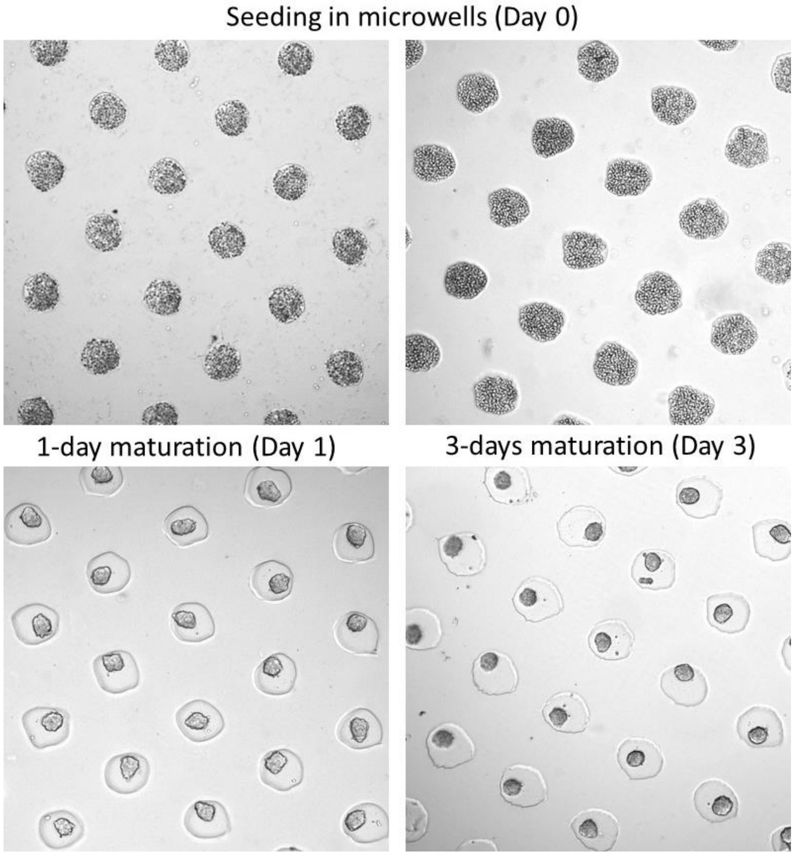


**Supplementary Figure 3**: Production of spheroids in agarose microwells from human mesenchymal stem cells. Optical microscopy images show cell aggregation and growth from initial cell seeding (day 0) up to 3 days of maturation into a spheroid configuration.

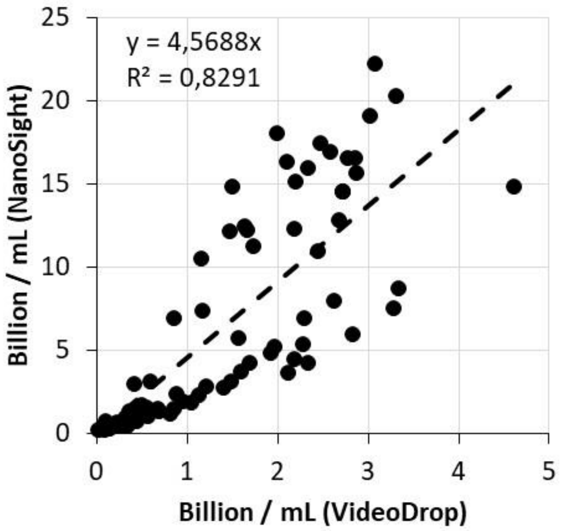


**Supplementary Figure 4**: Correlation between Nanoparticle Tracking Analysis (NTA) measurements performed with the NanoSight instrument and the VideoDrop instrument.

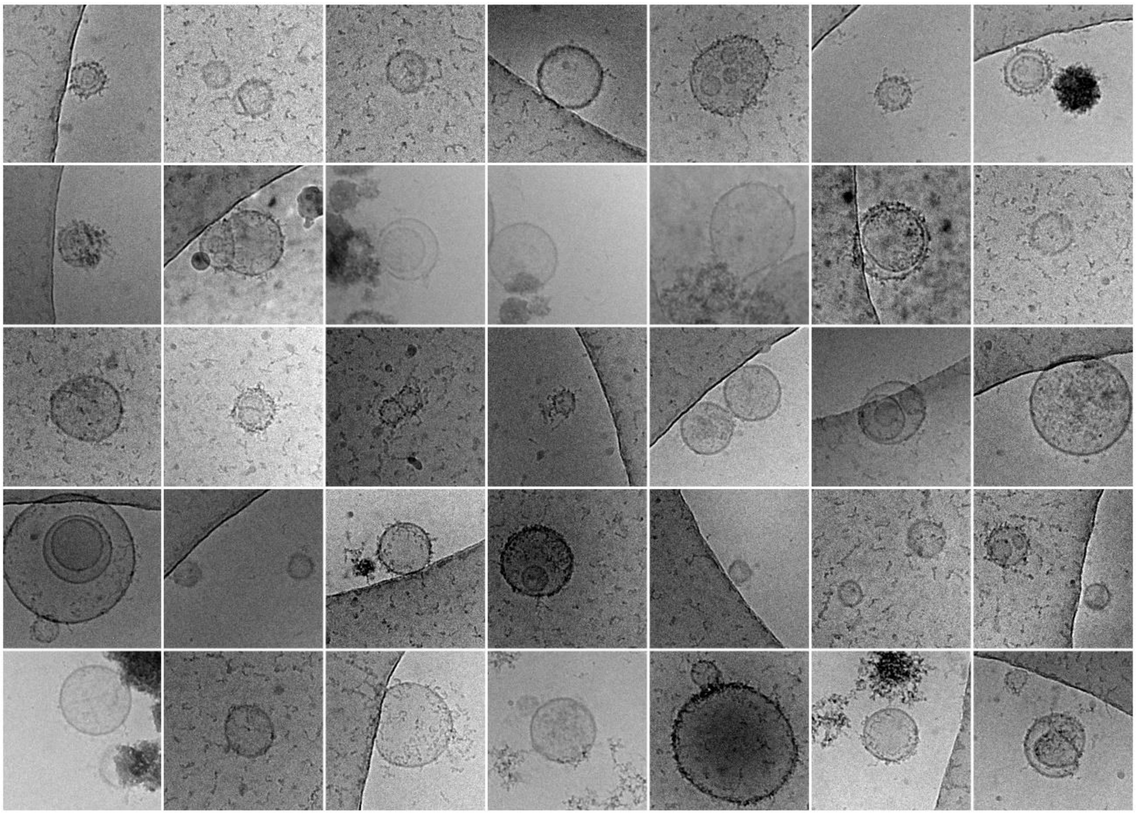

**Supplementary Figure 5**: CryoTEM observation of EVs produced under the Starv2D condition. Each square has a size of 300 nm x 300 nm.

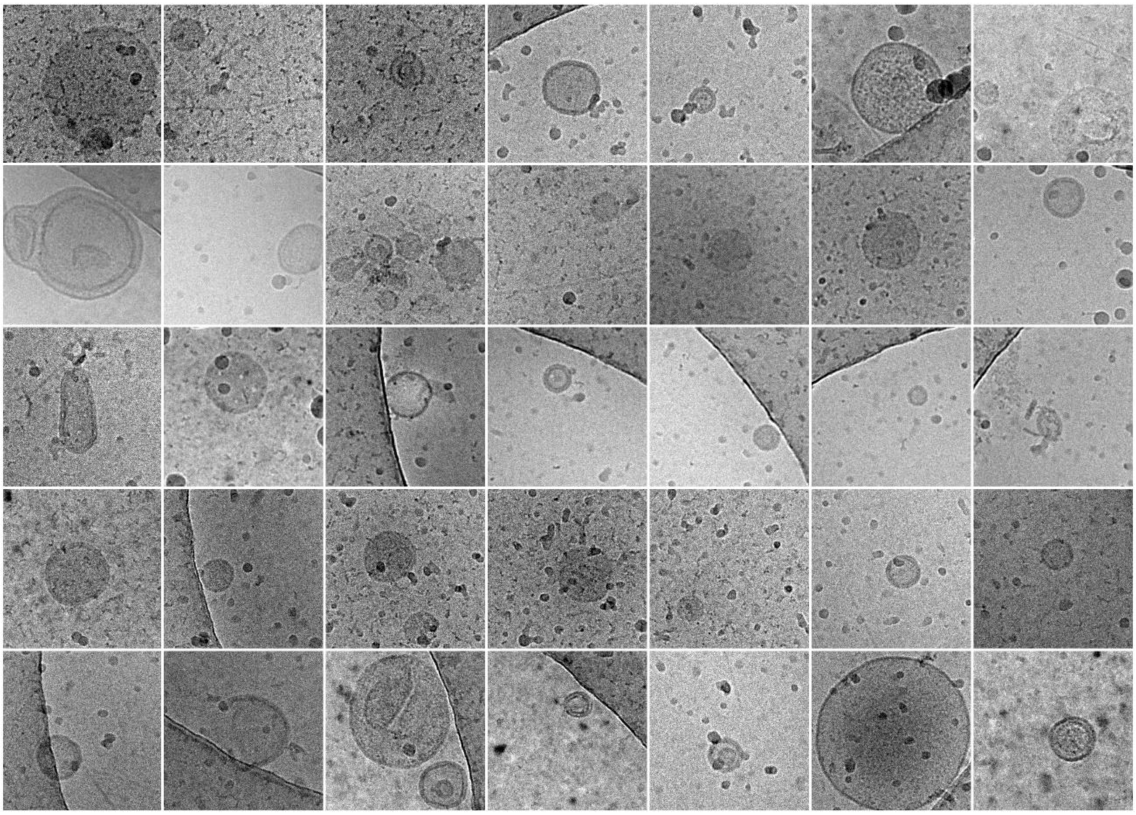

**Supplementary Figure 6**: CryoTEM observation of EVs produced under the Starv2D condition. Each square has a size of 300 nm x 300 nm.

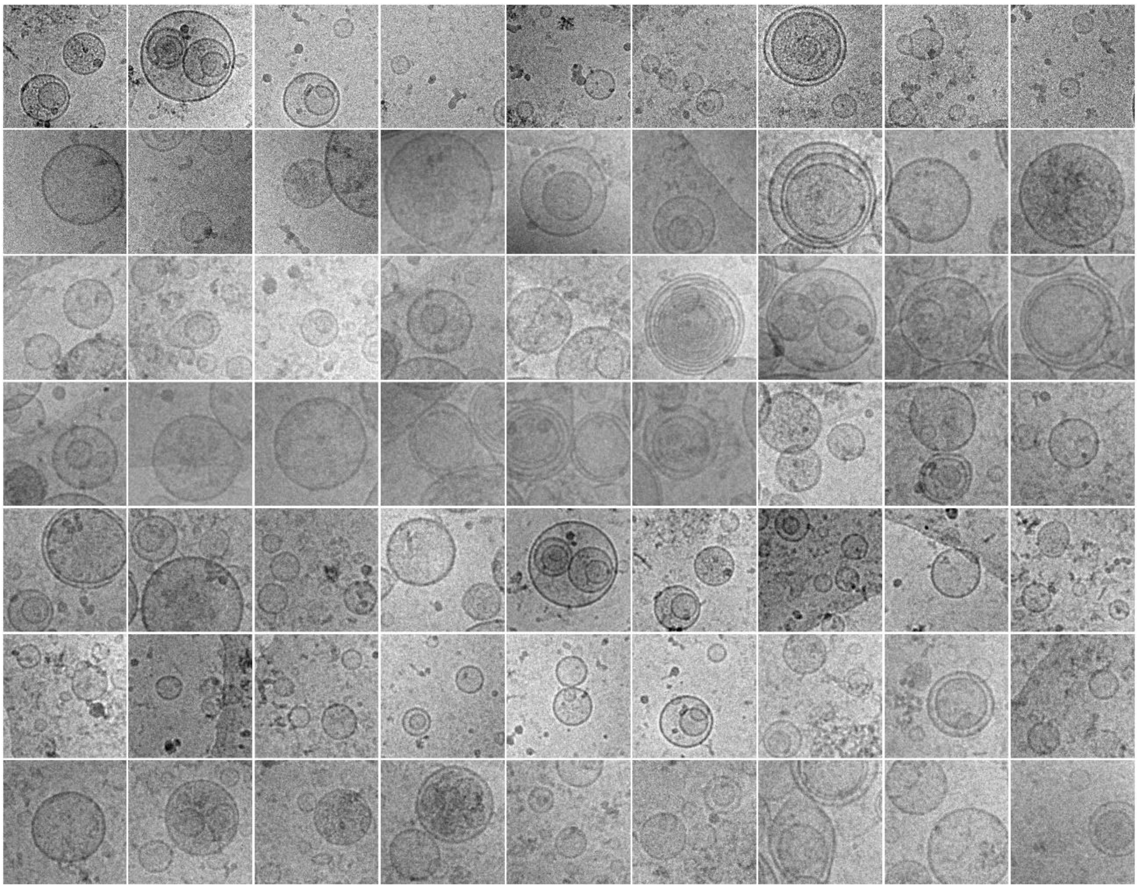

**Supplementary Figure 7**: CryoTEM observation of EVs produced under the HydroCell condition. Each square has a size of 300 nm x 300 nm.

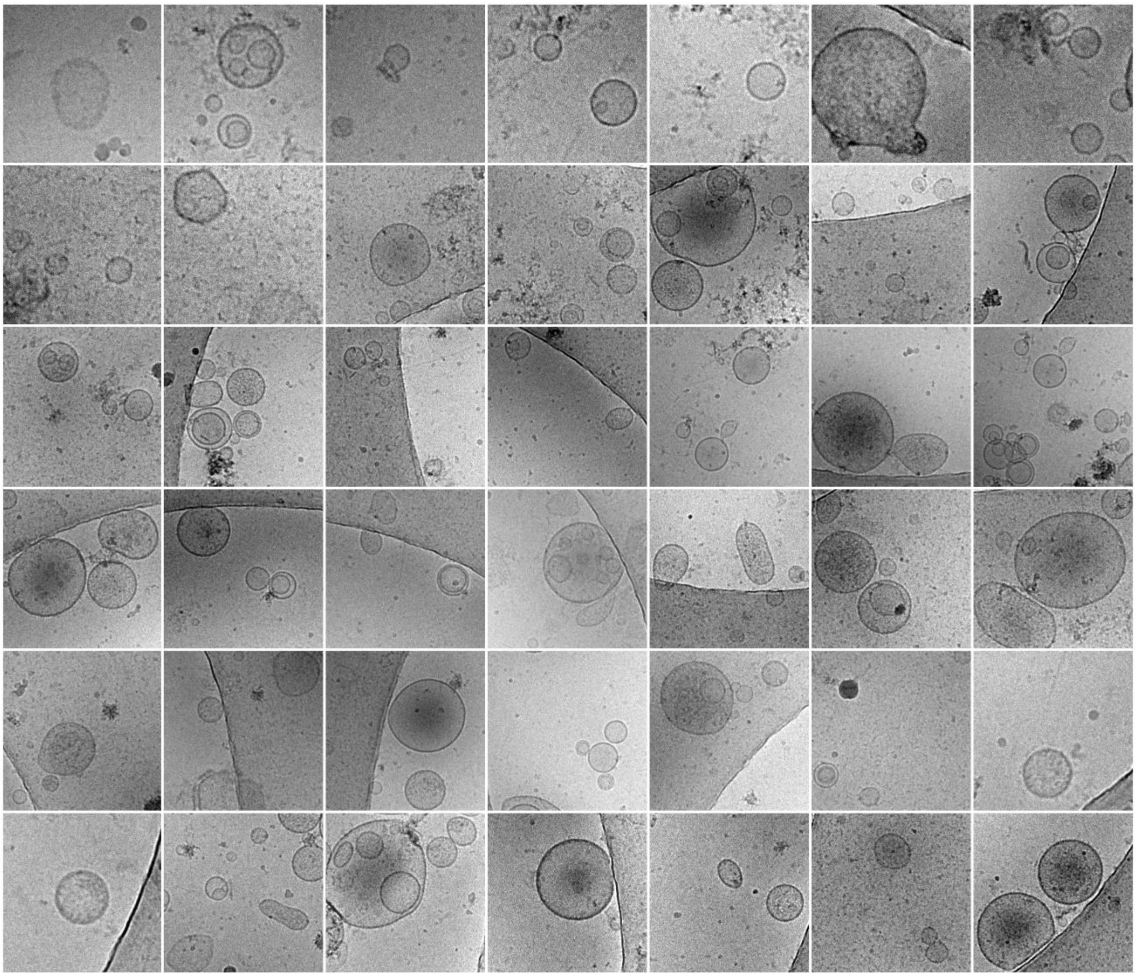

**Supplementary Figure 8**: CryoTEM observation of EVs produced under the HydroSph condition. Each square has a size of 300 nm x 300 nm.

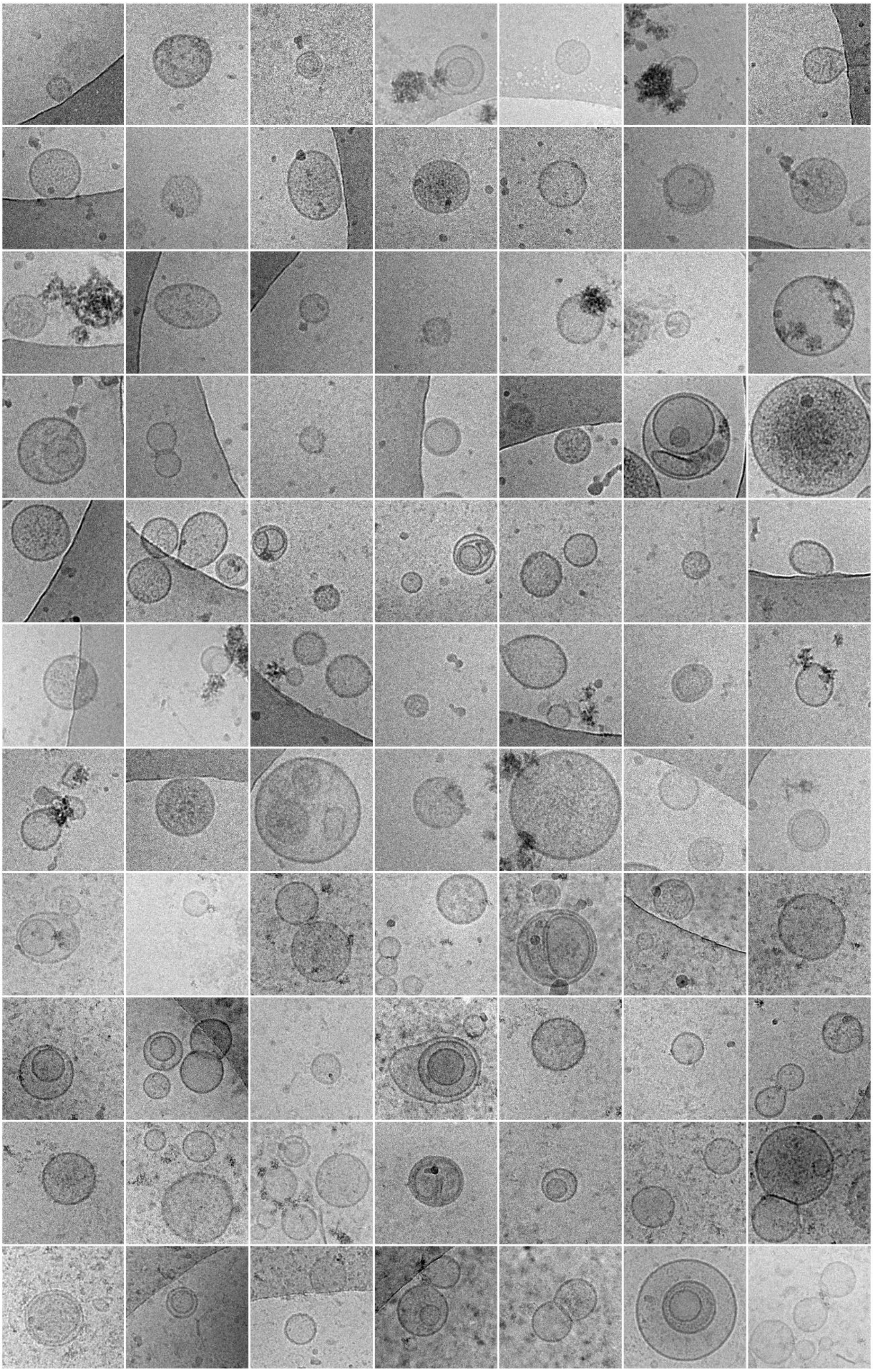

**Supplementary Figure 9**: CryoTEM observation of EVs produced under the HydroSphIn condition. Each square has a size of 300 nm x 300 nm.

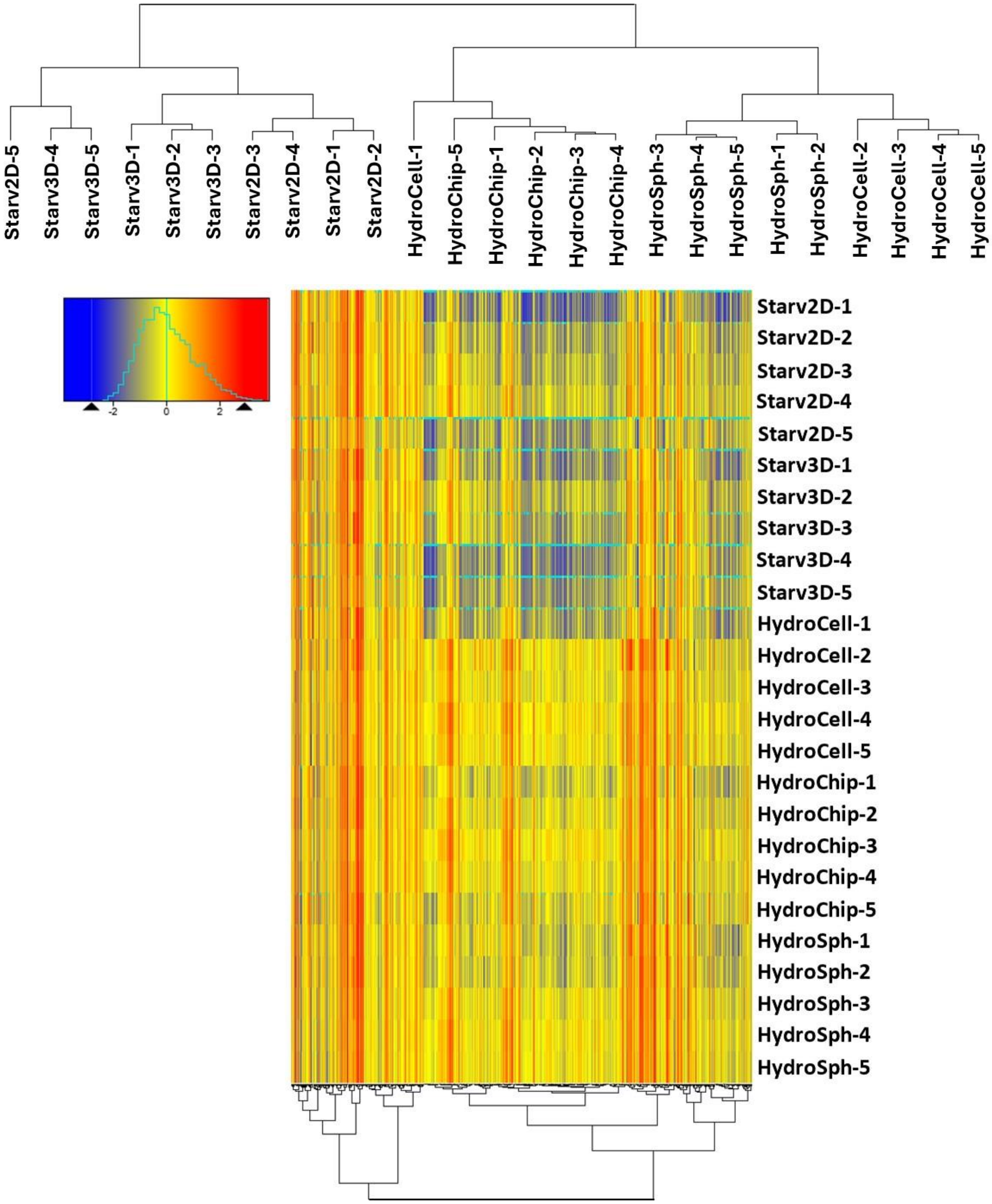


**Supplementary Figure 10**: Clustering of all proteins for all conditions.

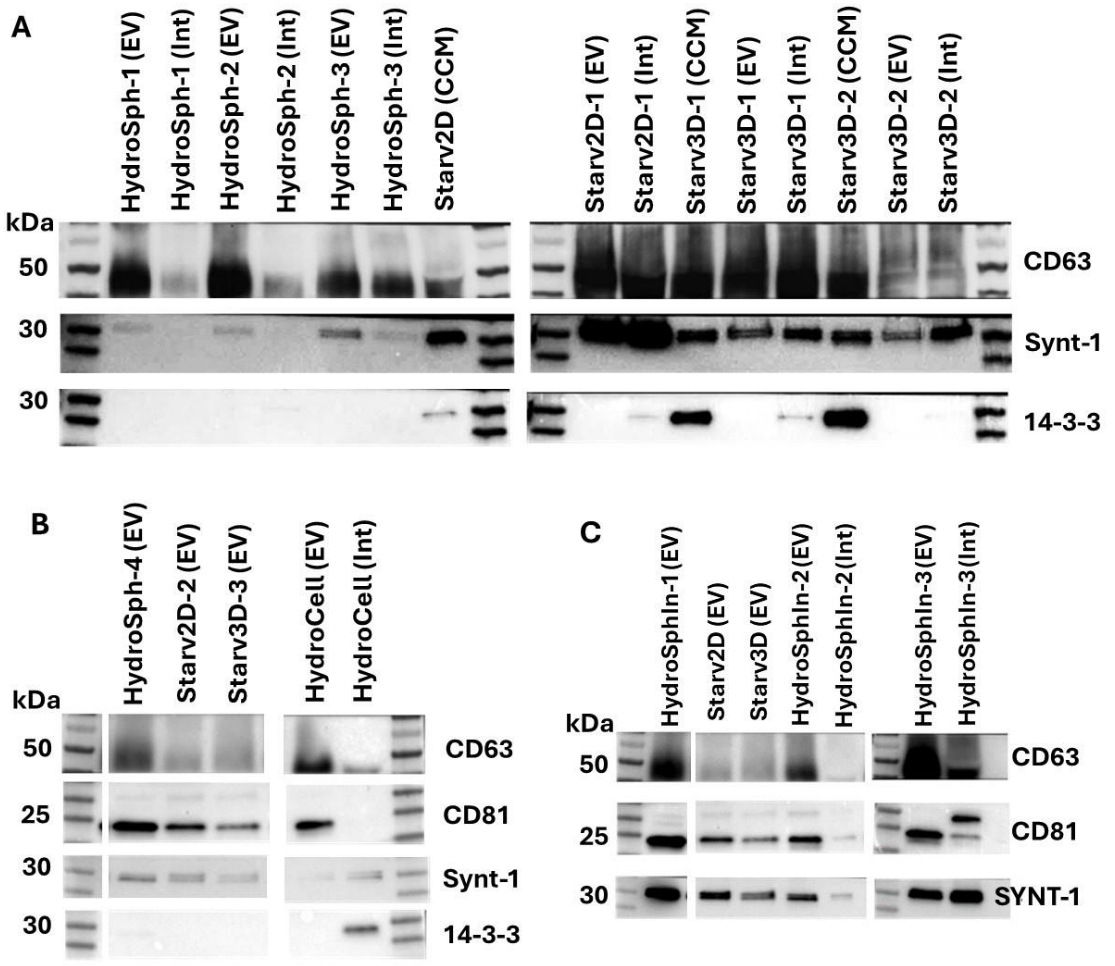


**Supplementary Figure 11**: **Western blot analysis**

***Method***: EVs samples were prepared for Western blot analysis by resuspending 30 µL of samples post SEC in 4x Laemmli Sample Buffer (Bio-Rad). The samples were then boiled for 10 minutes at 95°C and loaded onto 4–15% Mini-Protean® TGX Stain-Free™ gels (Bio-Rad) for electrophoretic separation under non-reducing conditions. Following electrophoresis, proteins were transferred onto Immuno-Blot PVDF membranes (Bio-Rad) using a semi-dry transfer method. After blocking, the membranes were incubated with primary anti-human antibodies: CD81 (clone TS81, Medix Biochema 1/1000), CD63 (clone H5C6, BD Bioscience 557305 1/1000), SDCBP (syntenin-1, clone EPR8102, Abcam 1/1000) and 14-3-3 (Clone EPR6380, Abcam, 1/1000). After primary antibody incubation, the membranes were post-incubated with horseradish peroxidase (HRP)-conjugated secondary antibodies goat anti-rabbit IgG (H + L) (Jackson 111-035-144) and goat anti-mouse IgG (H+L) (Jackson 111-035-146). After washing, immunoreactive bands were detected using Clarity western ECL substrate (Bio-Rad) with an exposure time of 16.250 seconds. The results were visualized using ChemiDoc™ Touch imager (Bio-Rad).

***A-B:*** Western blot analysis of specific EV membrane (CD63 and CD81) and cytosolic (Syntenin-1) markers, along with a co-isolated non-EV marker (14-3-3) for EVs and intermediate (Int) SEC fractions, or concentrated conditioned medium (CCM) produced from spheroids (HydroSph) or individual cells (HydroCell) in hydrodynamic conditions, and for EVs produced upon serum starvation from spheroids (Starv3D) or adherent cells (Starv2D). ***C:*** Western blots for the proteins CD63, CD81, and Synt-1, for EV fractions from control conditions Starv2D and Starv3D, and three independent HydroSphIn productions (HydroSphIn-1, -2, and -3). Intermediate fractions (Int) post SEC for conditions -2 and -3 are also shown.

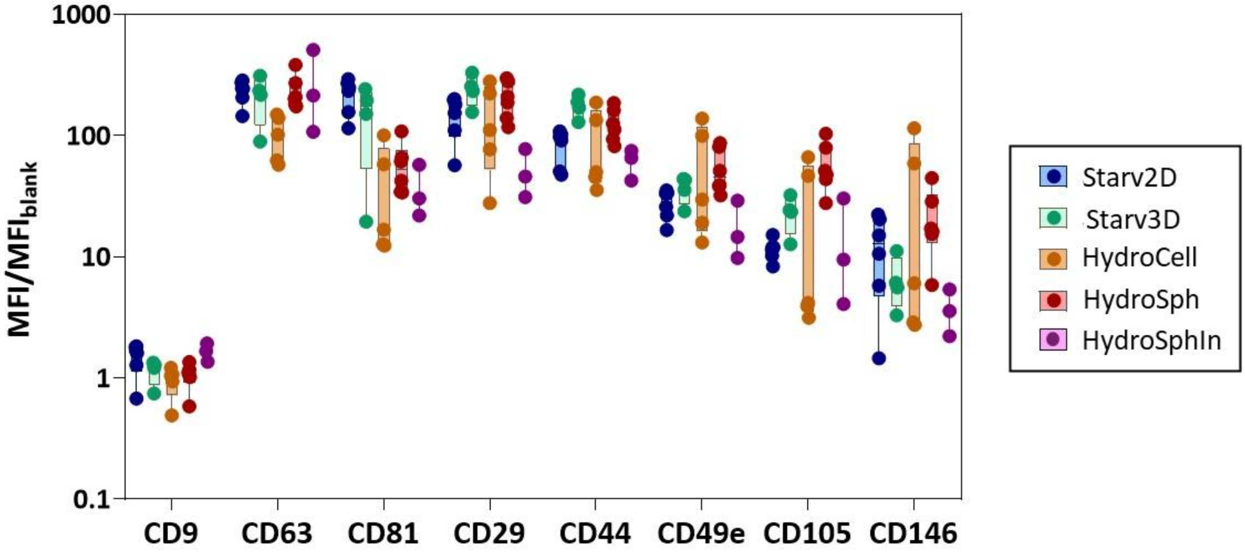


**Supplementary Figure 12**: **Bead-Based Multiplex Flow Cytometry Assay (MACSPlex)**

***Method***: EVs were analyzed using a bead-based multiplex assay for flow cytometry (MACSPlex EV Kit IO, human, Miltenyi) following the manufacturer's protocol. Briefly, EV concentration was quantified by nanoparticle tracking analysis (NTA), and the particle count was used to estimate the input EV quantity. A total of $5\times10^8$ EVs were diluted in MACSPlex buffer to achieve a final volume of 120 µL. Then, 15 µL of MACSPlex Exosome Capture Beads were added to the samples, which were incubated overnight at room temperature on an orbital shaker, shielded from light. After incubation, the samples were washed and then incubated with a mixture of APC-conjugated anti-CD9, anti-CD81, and anti-CD63 detection antibodies for 1 hour at room temperature. Following this, flow cytometry analysis was performed using an Aurora analyzer (Cytek). Data were processed and analyzed using FlowJo software (v10, FlowJo LLC). For data analysis, the 39 individual bead populations corresponding to different EV markers were gated, allowing the determination of the APC signal intensity for each respective bead type. Median fluorescence intensity (MFI) for each capture bead population was measured, and background signals were corrected by dividing the MFI values by the signal from non-EV controls, which underwent identical processing to the EV-containing samples.

***Graph****:* Bead-based fluorescence flow cytometric analysis of EV-specific and mesenchymal-specific protein markers in extracellular vesicles produced in hydrodynamic conditions from spheroids formed in agarose microwells (HydroSph) or formed in situ within the tubes (HydroSphIn) or from individual cells (HydroCell), and for EVs produced upon serum starvation from spheroids (Starv3D) or adherent cells (Starv2D). Results are presented as ratios of mean fluorescence intensity with respect to blank. Each point corresponds to an independent experiment.

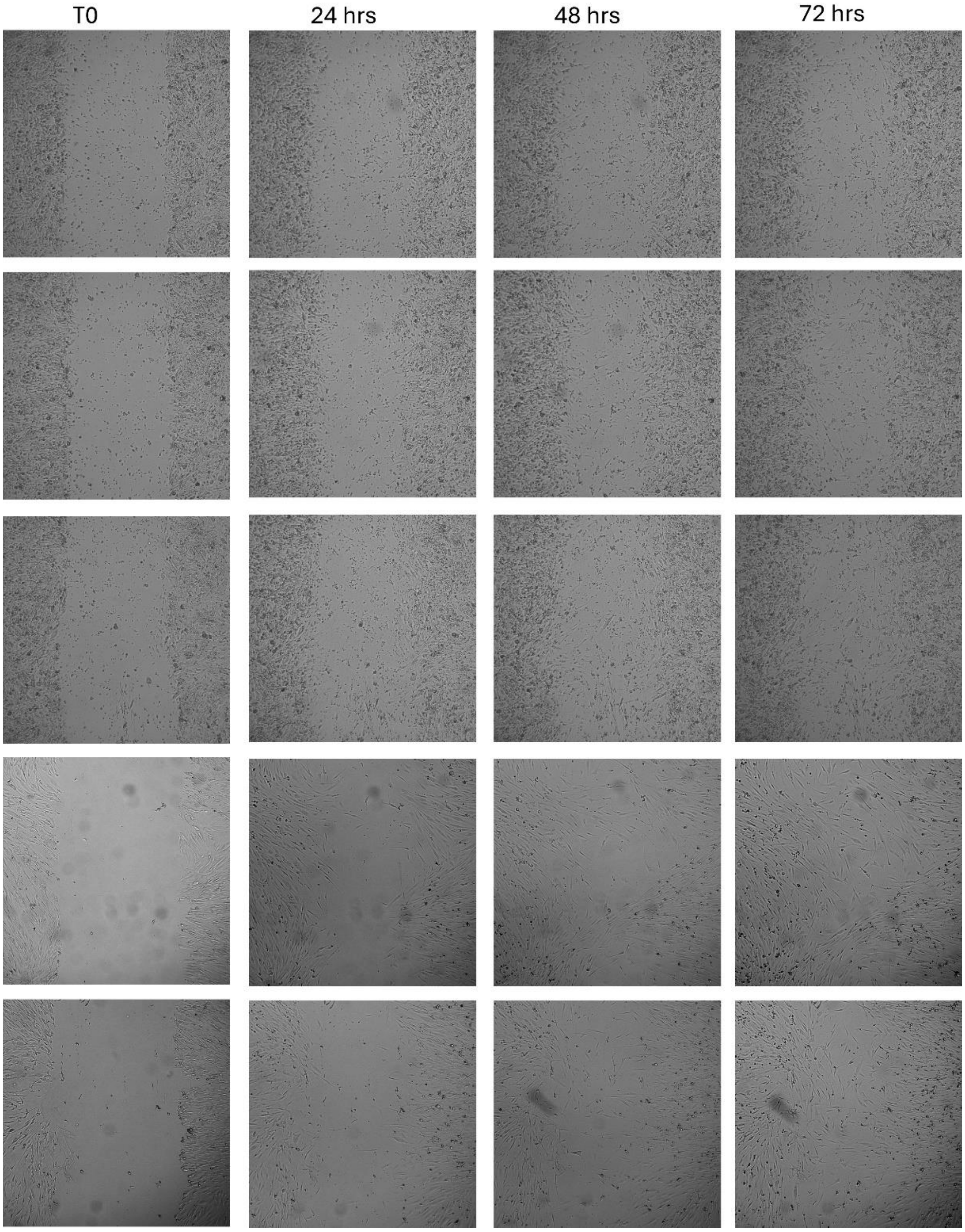


**Supplementary Figure 13A** : Images depicting the initial wound gap (T0) and wound healing at 24-hour, 48-hour and 72-hour time point, for the FBS 0% condition.

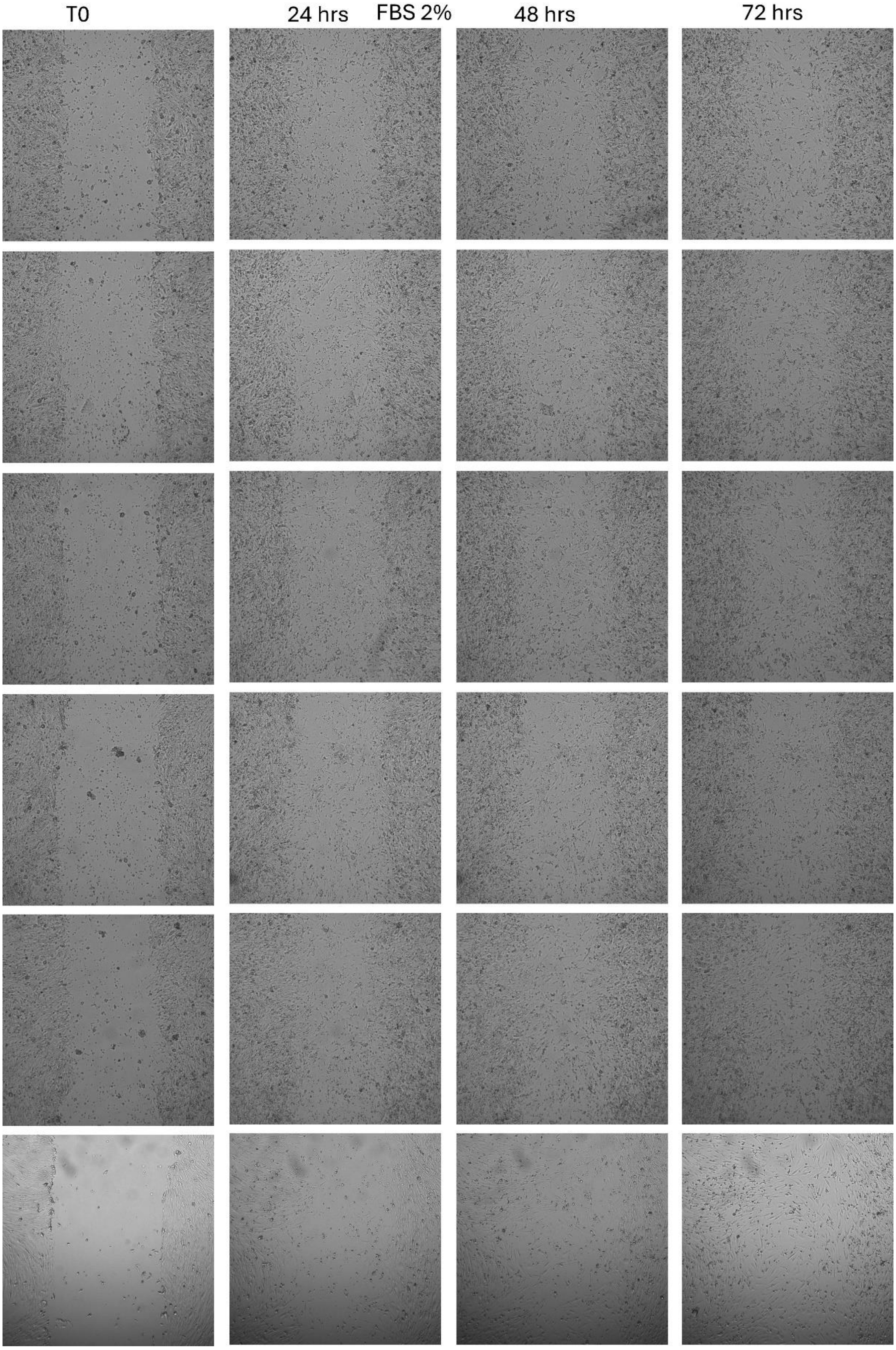


**Supplementary Figure 13B** : Images depicting the initial wound gap (T0) and wound healing at 24-hour, 48-hour and 72-hour time point, for the FBS 2% condition.

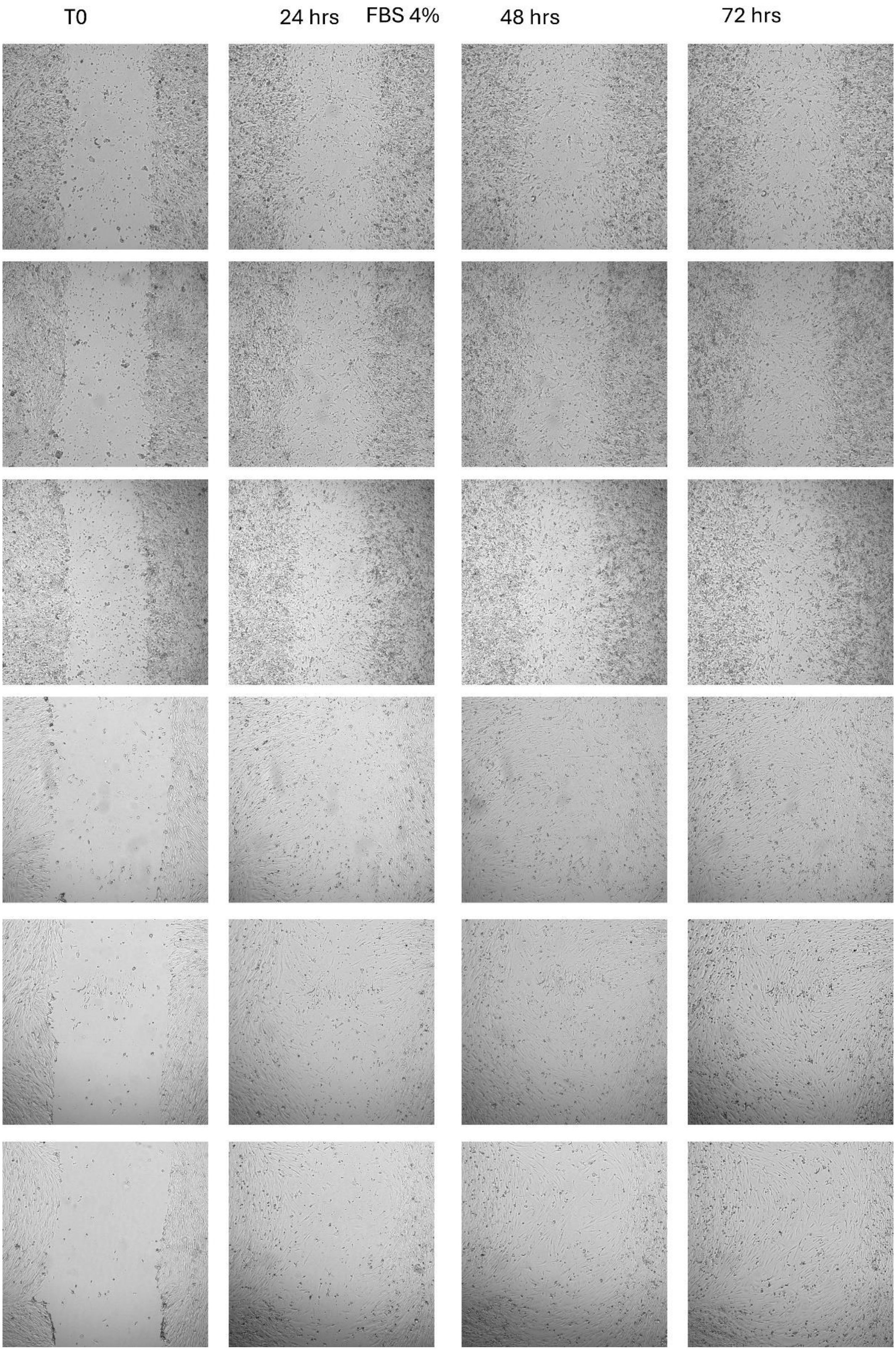


**Supplementary Figure 13C** : Images depicting the initial wound gap (T0) and wound healing at 24-hour, 48-hour and 72-hour time point, for the FBS 4% condition.

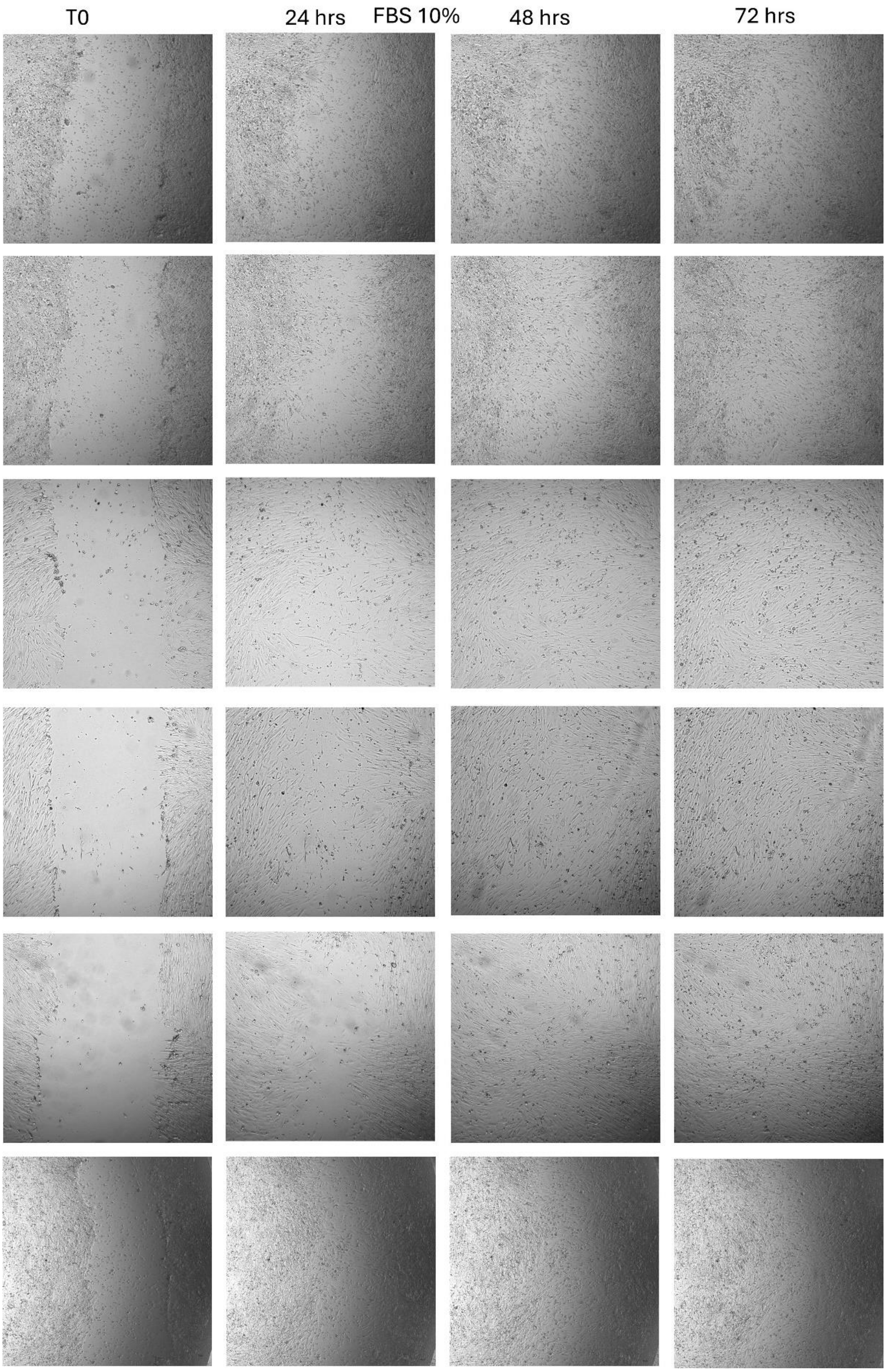


**Supplementary Figure 13D**: Images depicting the initial wound gap (T0) and wound healing at 24-hour, 48-hour and 72-hour time point, for the FBS 10% condition.

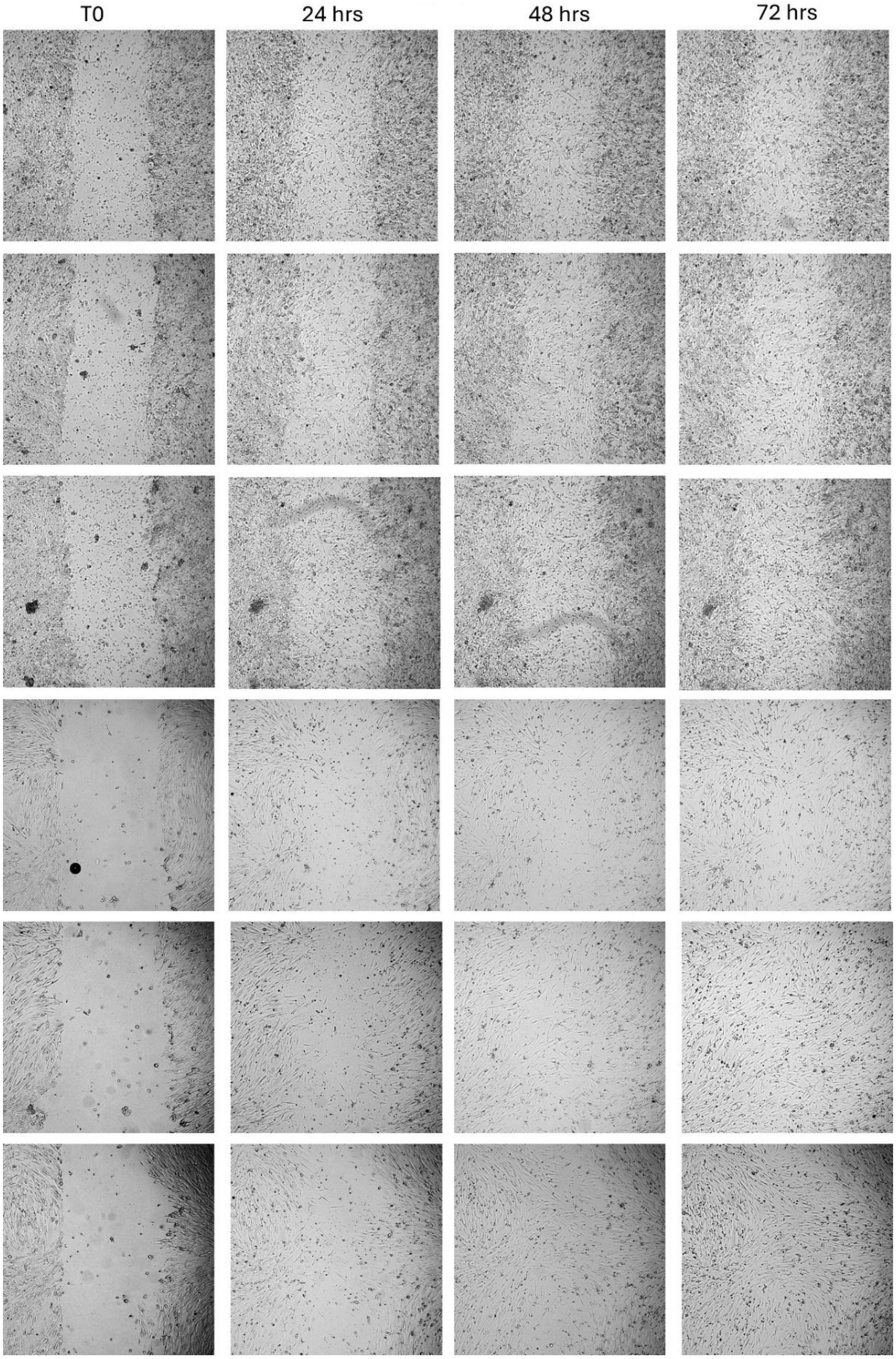


**Supplementary Figure 13E**: Images depicting the initial wound gap (T0) and wound healing at 24-hour, 48-hour and 72-hour time point, for the starvation 2D (Starv2D) condition.

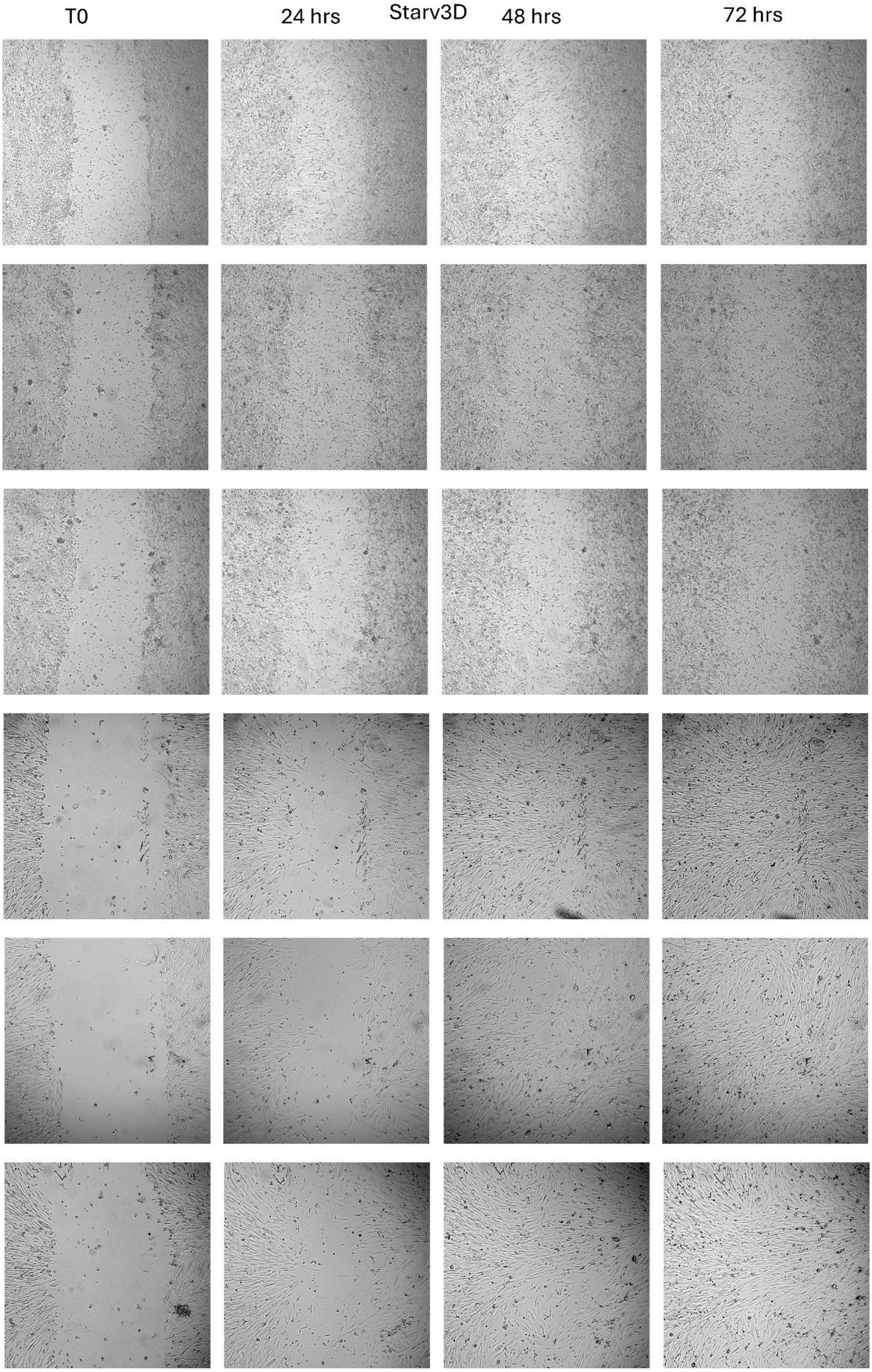


**Supplementary Figure 13F**: Images depicting the initial wound gap (T0) and wound healing at 24-hour, 48-hour and 72-hour time point, for the starvation 3D (Starv3D) condition.

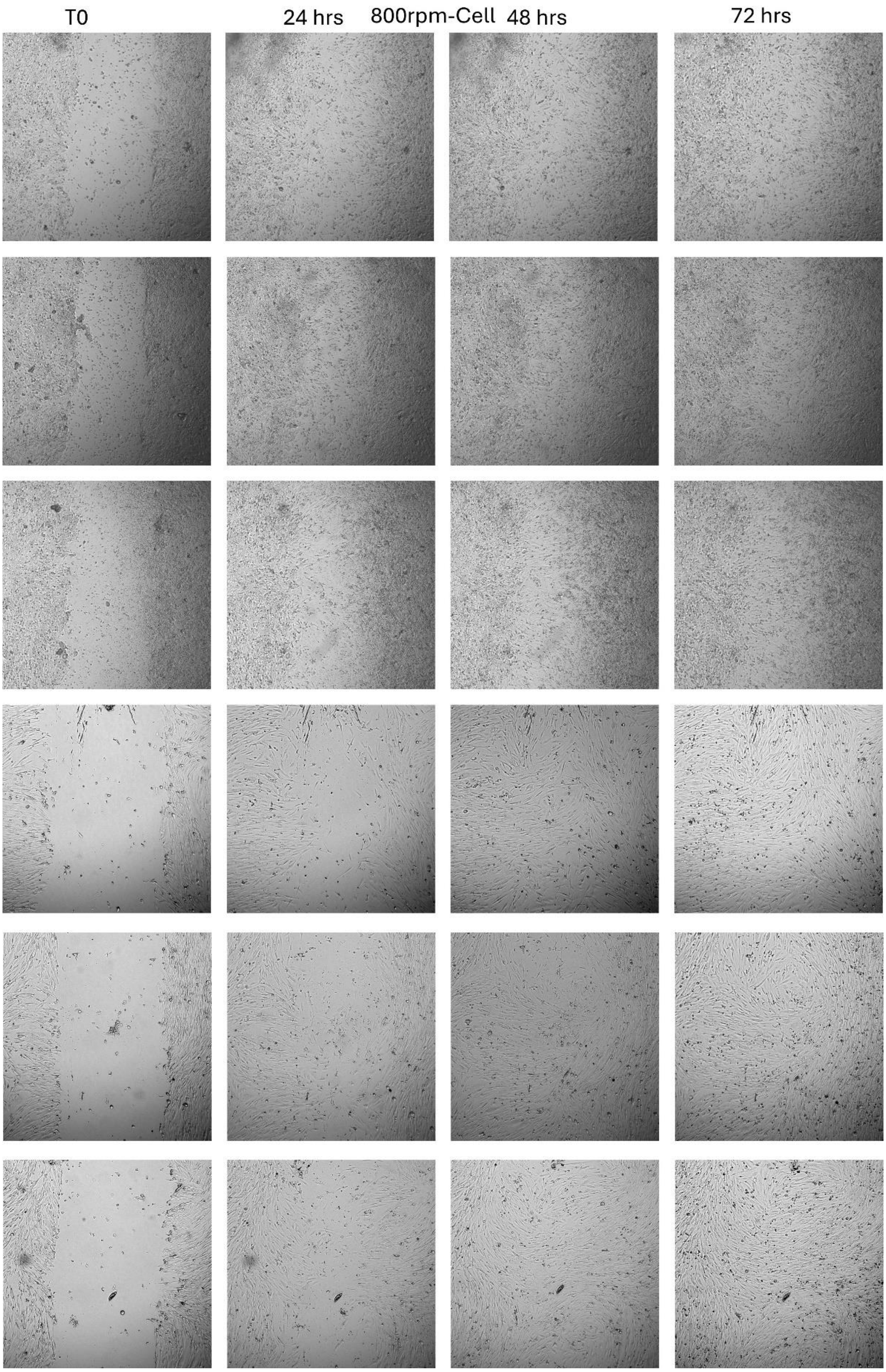


**Supplementary Figure 13G**: Images depicting the initial wound gap (T0) and wound healing at 24-hour, 48-hour and 72-hour time point, for the HydroCell condition.

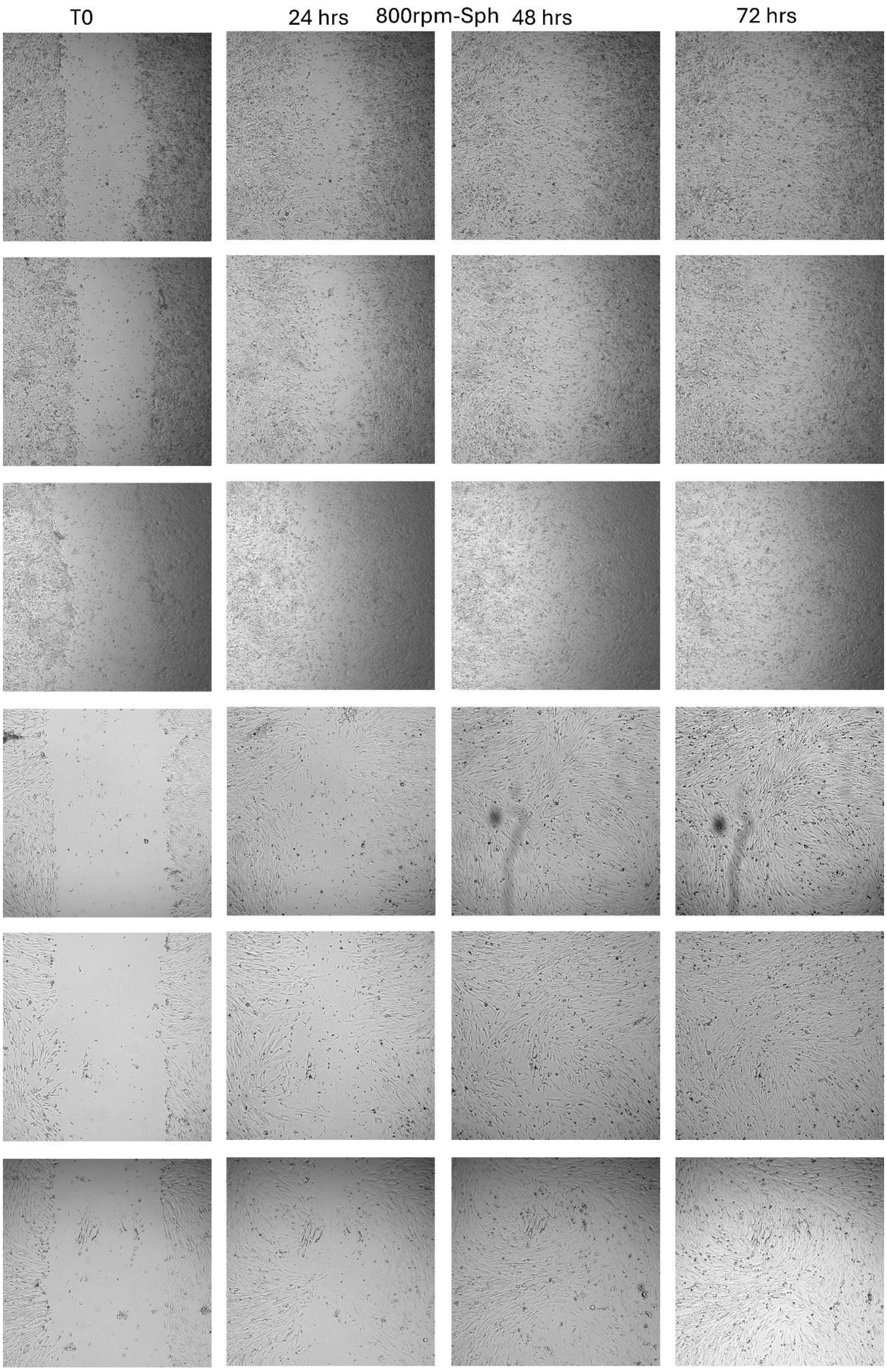


**Supplementary Figure 13H**: Images depicting the initial wound gap (T0) and wound healing at 24-hour, 48-hour and 72-hour time point, for the HydroSph condition.

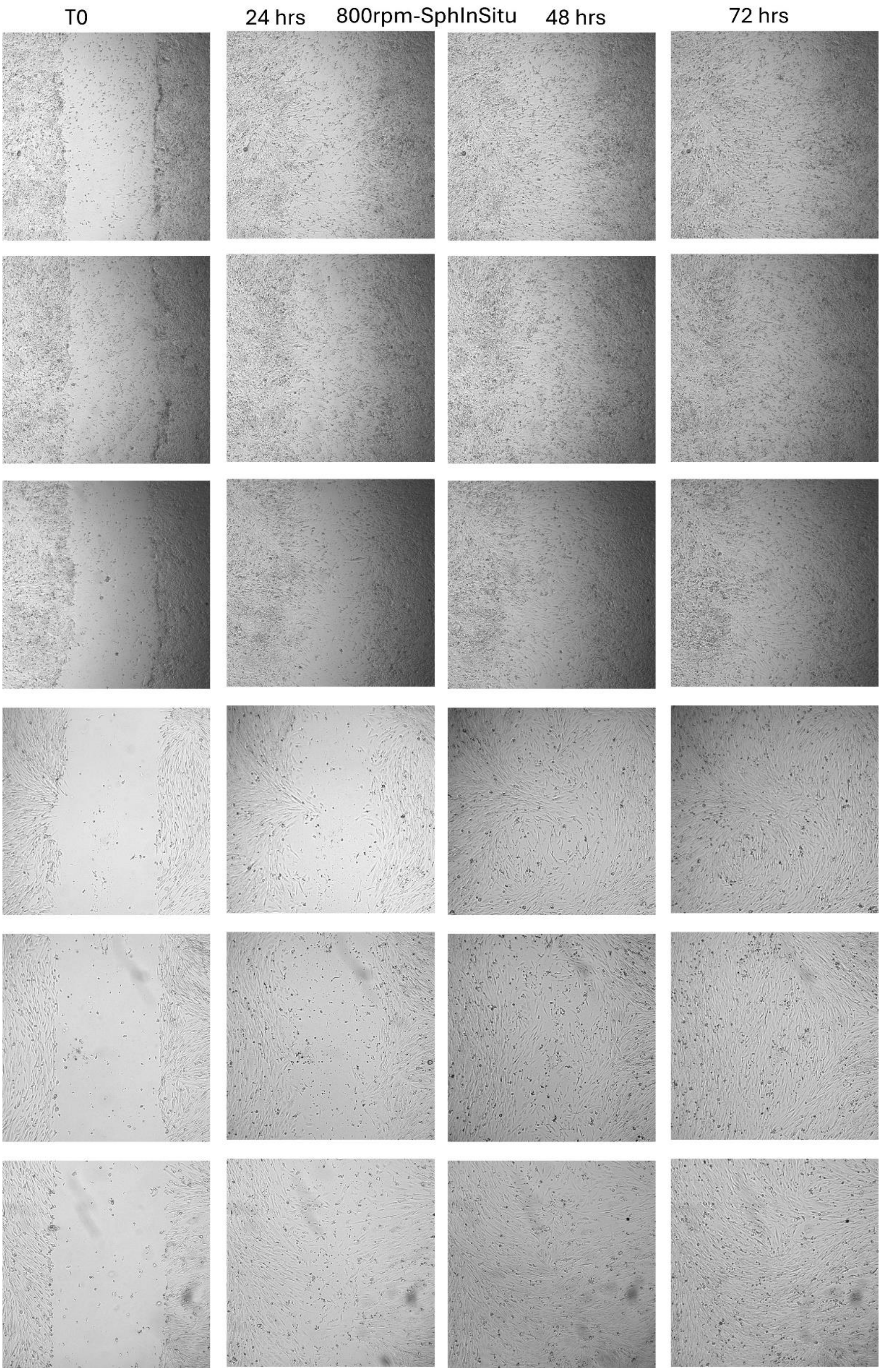


**Supplementary Figure 13I**: Images depicting the initial wound gap (T0) and wound healing at 24-hour, 48-hour and 72-hour time point, for the HydroSphIn condition.

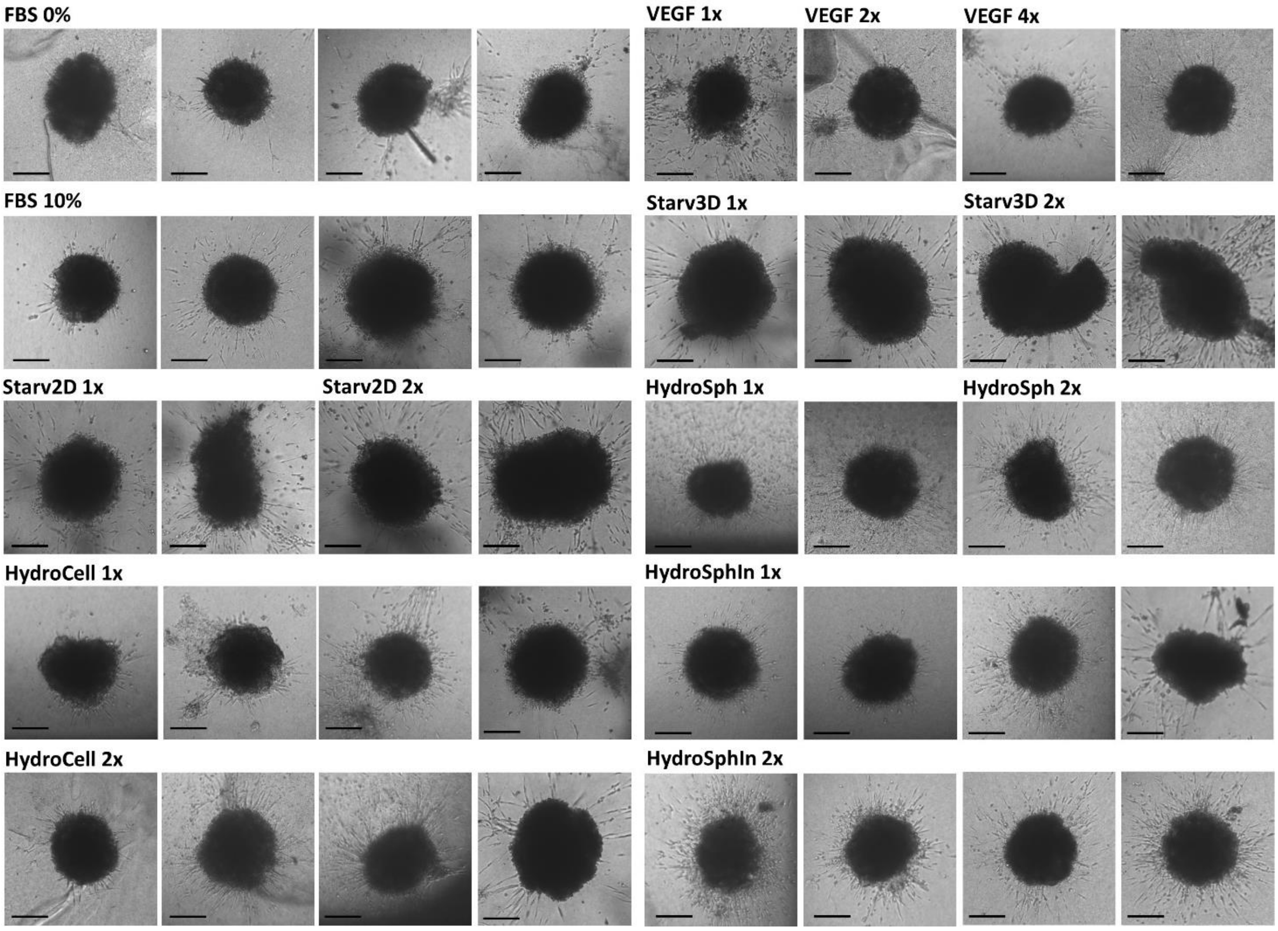


**Supplementary Figure 14**: Images of sprout formation from endothelial spheroids, illustrating the neo-angiogenesis capacity, for all conditions depicted in Fig.3c and Fig.5d. Scale bars = 100 µm.

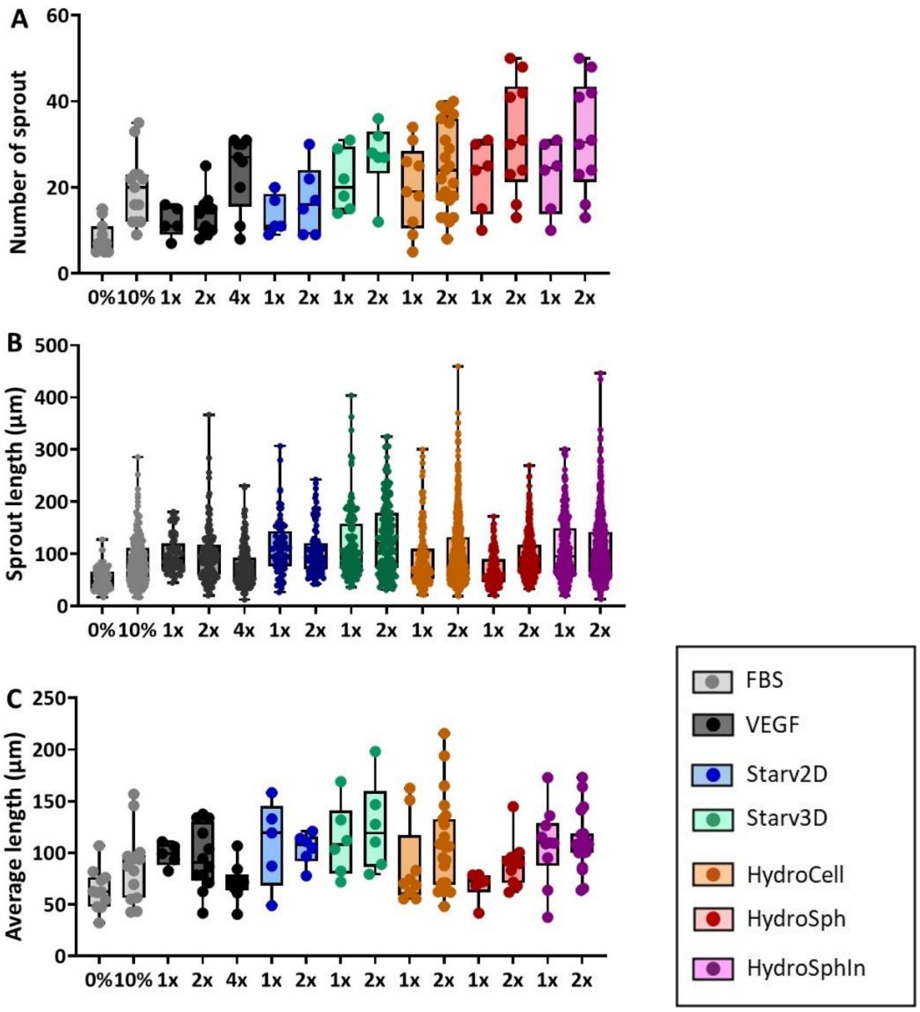


**Supplementary Figure 15**

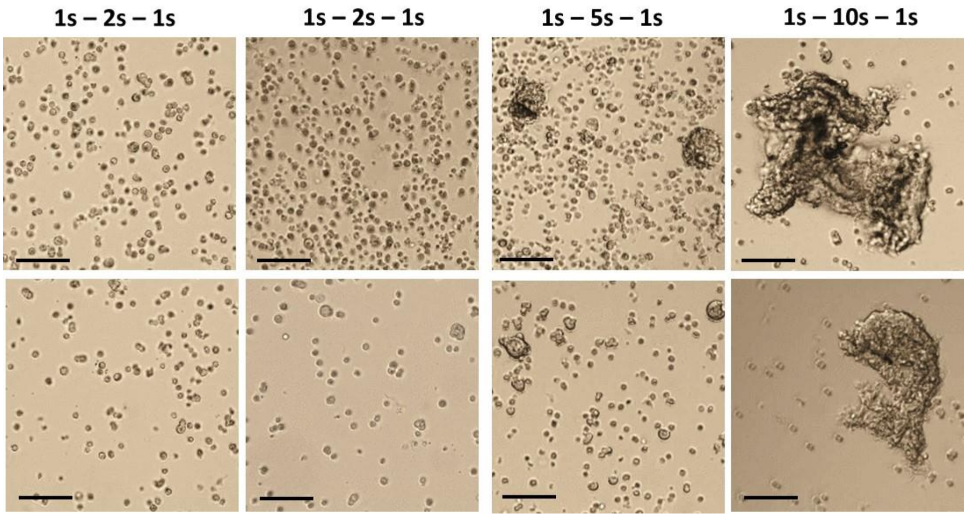


**Supplementary Figure 16A.** Scale bars = 200 µm.

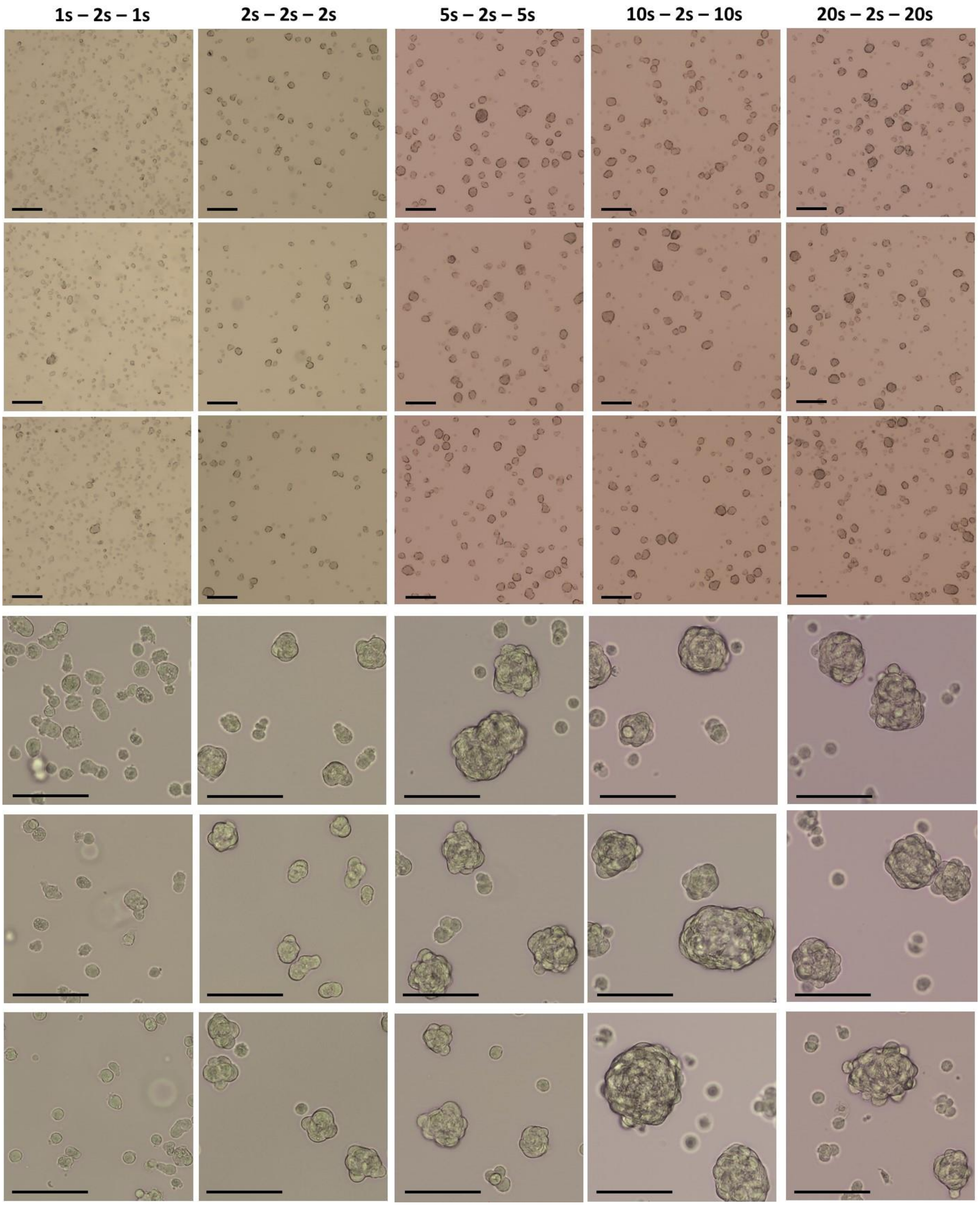


**Supplementary Figure 16B.** Scale bars = 200 μm.

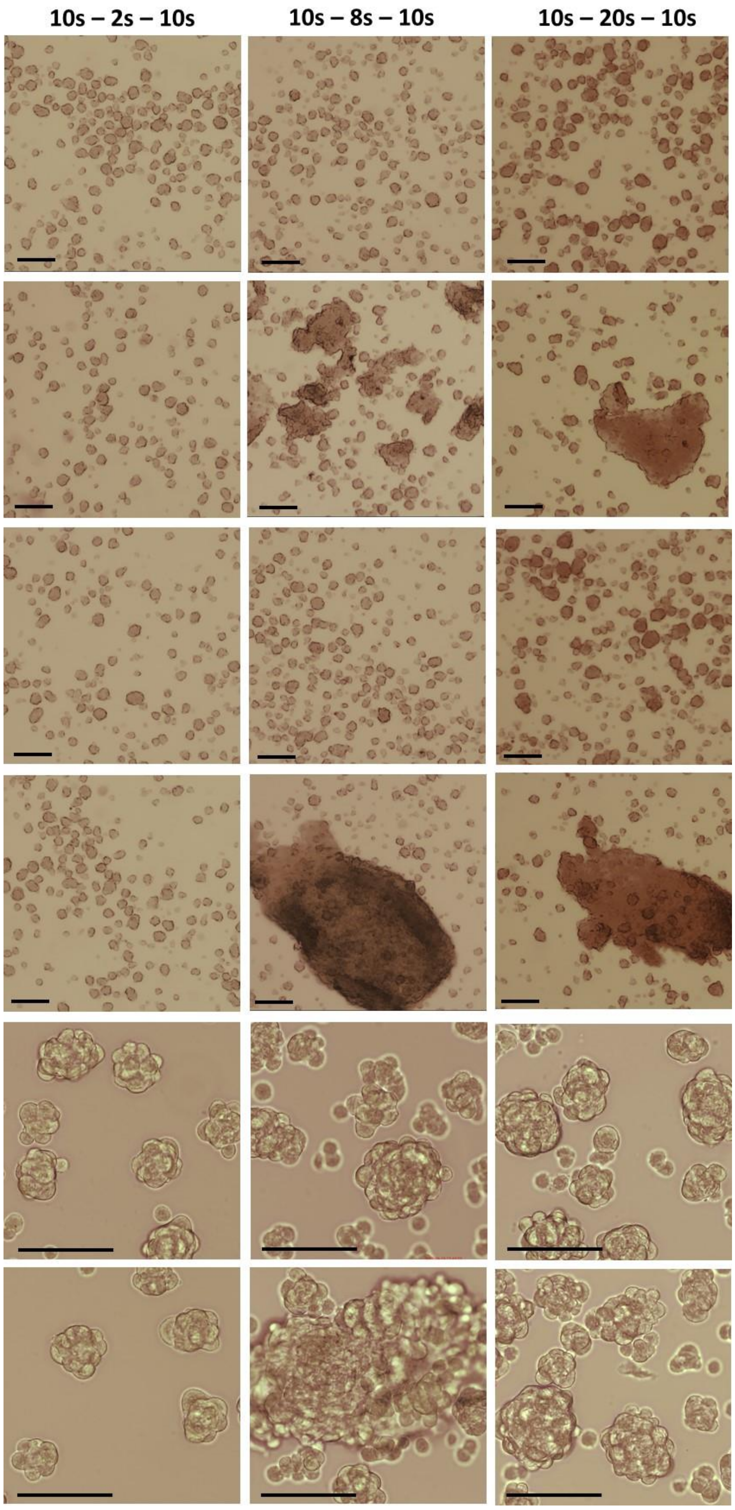


**Supplementary Figure 16C.** Scale bars = 200 µm.

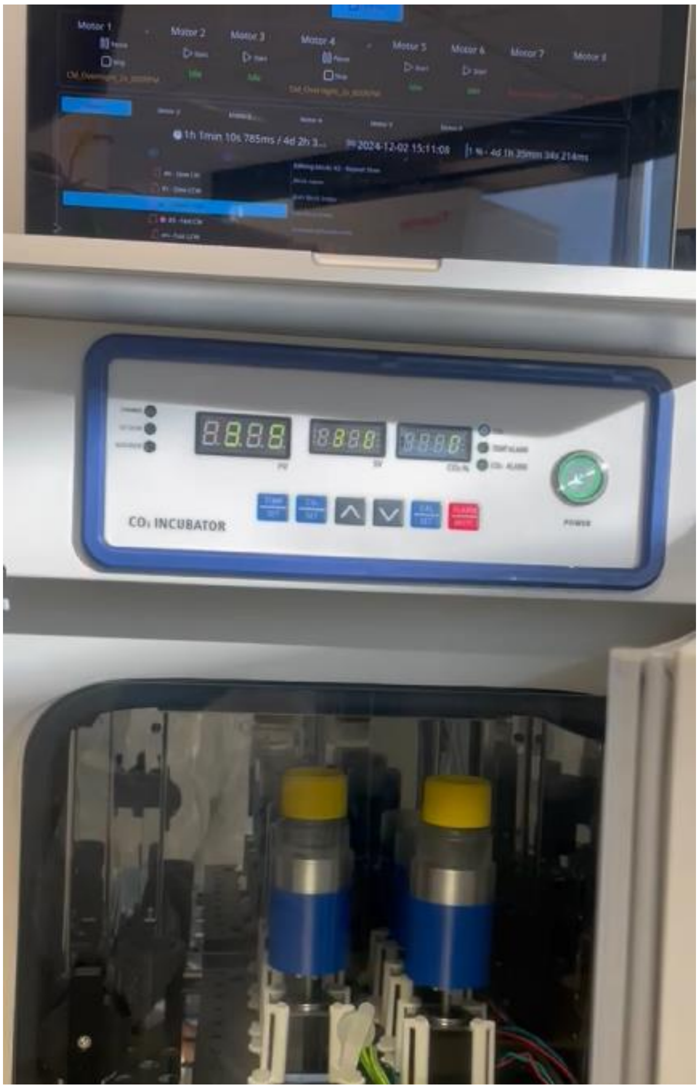


**Supplementary Figure 17**: Prototype.

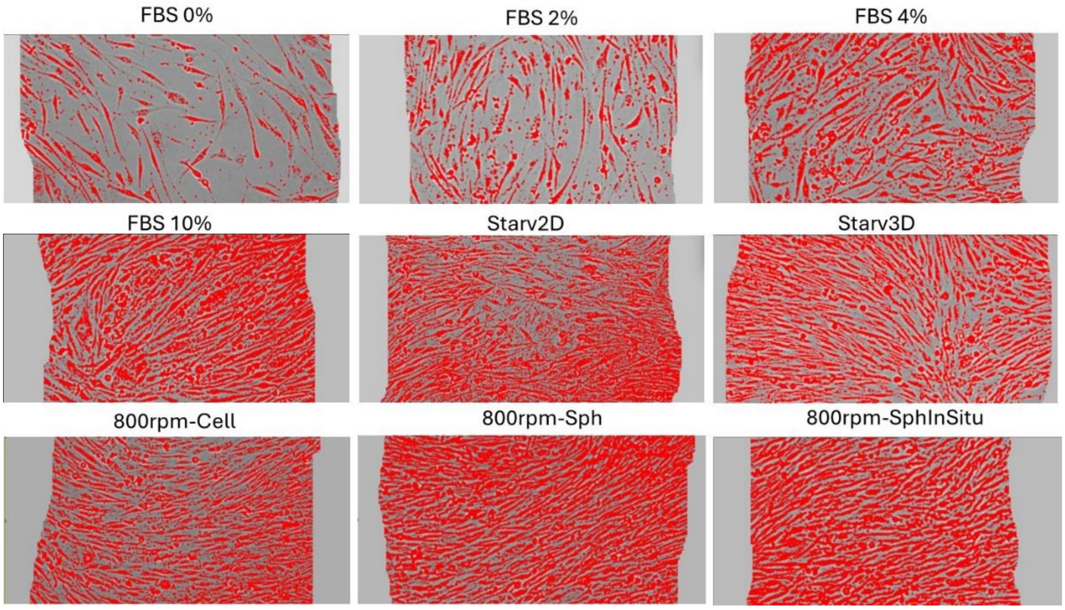


**Supplementary Figure 18**: Example of thresholding for wound healing.
